# Proximal and Distal Nephron-specific Adaptation to Furosemide

**DOI:** 10.1101/2021.01.12.426306

**Authors:** Aram J. Krauson, Steven Schaffert, Elisabeth M. Walczak, Jonathan M. Nizar, Gwen M. Holdgate, Sonali Iyer, Ragwa Elsayed, Alexandre Gaudet, Purvesh Khatri, Vivek Bhalla

**Affiliations:** Division of Nephrology, Department of Medicine, Stanford University School of Medicine, Stanford, CA; Division of Biomedical Informatics Research, Department of Medicine, Stanford University School of Medicine, Stanford, CA; BD Biosciences

**Keywords:** diuretic, tubular remodeling, diuretic resistance, ion transport, insulin-like growth factor 1, hyperplasia, hypertrophy

## Abstract

Furosemide, a widely prescribed diuretic for edema-forming states, inhibits sodium reabsorption in the thick ascending limb of the nephron. Tubular adaptation to diuretics has been observed, but the range of mechanisms along the nephron has not been fully explored. Using morphometry, we show that furosemide induces renal tubular epithelial hyperplasia selectively in distal nephron segments. By comparison, we find progressive cellular hypertrophy in proximal and distal nephron segments. We next utilize single cell RNA sequencing of vehicle- and furosemide-treated mice to define potential mechanisms of diuretic resistance. Consistent with distal tubular cell hyperplasia, we detect a net increase in DCT cell number and *Birc5*, an anti-apoptotic and pro-growth gene, in a subset of DCT cells, as the most prominently up-regulated gene across the nephron. We also map a gradient of cell-specific transcriptional changes congruent with enhanced distal sodium transport. Furosemide stimulates expression of the mitogen IGF-1. Thus, we developed a mouse model of inducible deletion of renal tubular IGF-1 receptor and show reduced kidney growth and proximal, but not distal, tubular hypertrophy by furosemide. Moreover, genes that promote enhanced bioavailability of IGF-1 including *Igfbp1* and *Igfbp5* are significantly and differentially expressed in proximal tubular segments and correspond to IGF-1R-dependent hypertrophy. In contrast, downstream PI3-kinase signaling genes including *Pdk1, Akt1, Foxo3, FKBP4, Eif2BP4*, and *Spp1* are significantly and differentially expressed in distal nephron segments and correspond to IGF-1R-independent hypertrophy. These findings highlight novel mechanisms of tubular remodeling and diuretic resistance, provide a repository of transcriptional responses to a common drug, and expand the implications of long-term loop diuretic use for human disease.

## INTRODUCTION

An estimated five million Americans require long-term treatment for edema due to congestive heart failure, cirrhosis, nephrotic syndrome, or chronic kidney disease^1-4^. Furosemide, a common treatment for these edema-forming states, increases urine sodium excretion by inhibition of the Na:K:2Cl cotransporter (NKCC2) in renal tubular epithelial cells of the thick ascending limb (TAL) of the loop of Henle^5^. This nephron segment is responsible for 20 – 30% of the filtered sodium reabsorbed^6,7^ **[Figure 1a]**. Furosemide and other “loop” diuretics also alter the balance of other plasma electrolytes with implications for neuronal and cardiomyocyte function.

**Figure 1:**
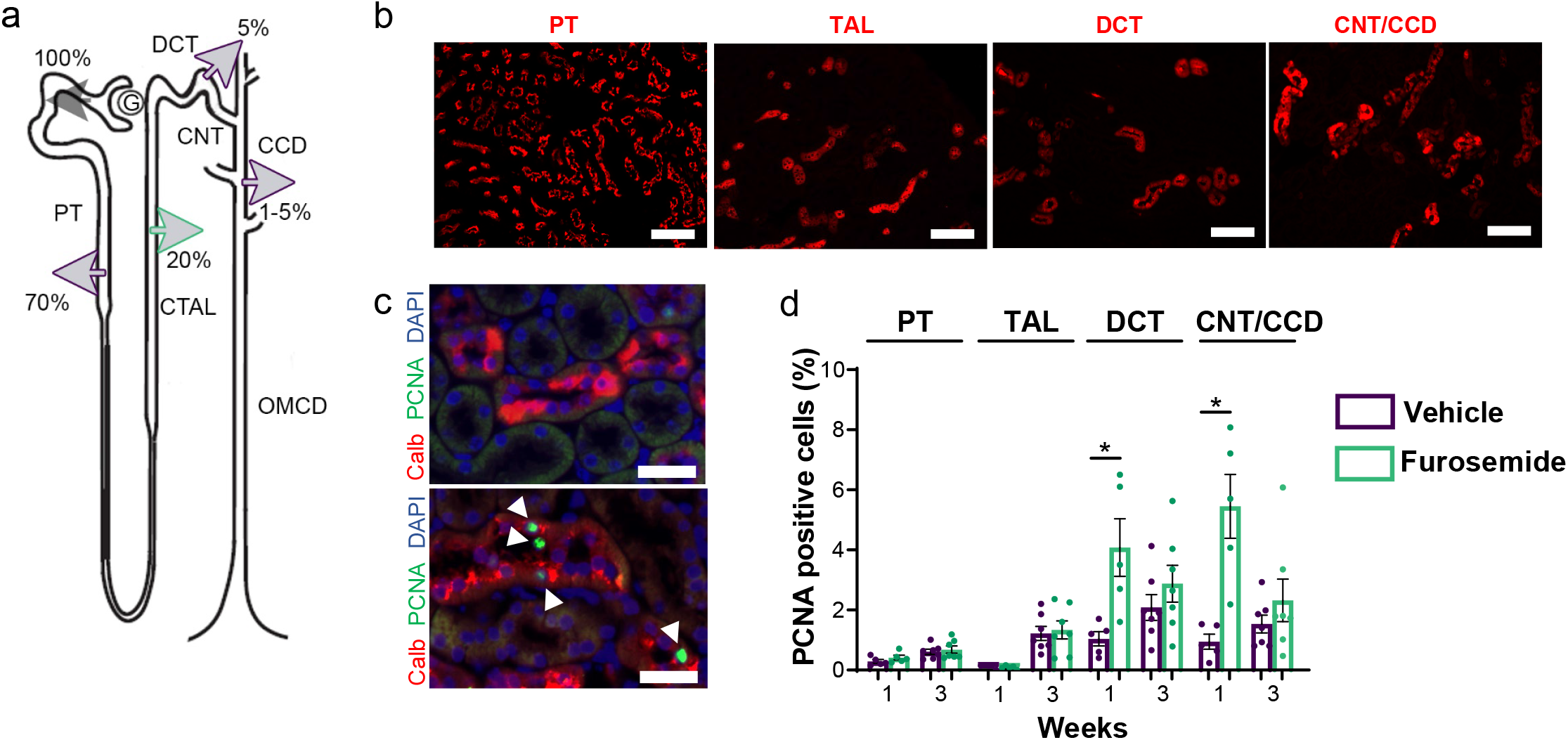
One-week furosemide treatment induces hyperplasia in the distal nephron. (a) Localization of furosemide action (*green*) along the nephron relative to % of sodium reabsorption. (b) Nephron-specific markers for the PT, TAL, DCT, and CNT/CCD segments (*red*) (right panels). Scale bars: 100 µm. (c) Sample images of vehicle-(top) and furosemidetreated mice (bottom) show calbindin (*red*) in combination of PCNA (*green*) and DAPI (*blue*) for cell proliferation analysis. Overlapping fluorescence of PCNA with nuclei staining (white arrowheads) within segments were counted. Scale bars: 50 µm. (d) Histogram of PCNA positive cell percentages (mean ±SEM) comparing vehicle- and furosemide-treated mice at 1 and 3 weeks (N=5-7 mice/group). *p-value < 0.05 versus vehicle treatment.

Resistance to loop diuretics is defined as an increase in sodium reabsorption in nephron segments distal to the loop of Henle. Diuretic resistance mandates the need for higher doses and/or additional classes of diuretics, some with potential adverse side effects; diuretic resistance is also associated with mortality in heart failure and other edematous states^8-13^. While the nephron is known to be the primary site of diuretic resistance^14^, the mechanisms are unclear. Tubular adaptation to diuretics can be due to an increase activity/expression of sodium transporters, e.g. the Na-Cl cotransporter (NCC, *Slc12a3*) or the epithelial sodium channel (ENaC, *Scnn1a,1b, 1g*)^15,16^. Another source of adaptation is structural remodeling of these segments via cell proliferation (hyperplasia), cell growth (hypertrophy)^9,17,18^, loss of segments through cell atrophy or death^19^ or transdifferentiation through reprogramming.

Changes in sodium transport within a particular segment may correlate with tubular epithelial cell plasticity along the nephron. For example, with inhibition of NCC, NCC-expressing cells in the distal convoluted tubule (DCT) show apoptosis^19^ whereas increased activation of NCC increases the mass of DCT^20,21^. Furosemide treatment increases the kidney-to-body weight ratio in rodent studies^22,23^. The mechanisms of cell plasticity are unknown and hypertrophy of only a short segment such as the DCT is unlikely to account for increases in weight of the total kidney. Thus, we postulate that there may be diverse changes in gene expression and/or nephron architecture over time along the tubule responsible for the increase in kidney mass and diuretic resistance, beyond the previously observed distal convoluted tubule and collecting duct hypertrophy^15^.

Diuretic-induced tubular modeling has been previously associated with renal insulin-like growth factor-1 (IGF-1) signaling^24^. IGF-1 is a secreted growth factor that binds the receptor tyrosine kinase IGF-1R, to promote proliferation, cell growth, and survival ^24-26^. Mitogenic IGF-1 and IGF-1 binding proteins accumulate in the kidney proceeding hypertrophy^27^, and induction of IGF-1 overexpression in untreated tubular epithelial cells induces hypertrophy^28,29 30^. However, whether IGF-1R signaling directly contributes to diuretic-induced hypertrophy is unknown.

In this study we examine the diversity of cell-type specific mechanisms of furosemide-induced tubular remodeling using tubular morphometry and single cell RNA sequencing (RNAseq).

## RESULTS

### Furosemide induces distal tubular cell-specific hyperplasia

We examined changes in tubular epithelial cell proliferation of vehicle- and furosemide-treated mice over time [**Figure S1**] in four representative nephron segments (PT, TAL, DCT, and principal cells of the connecting tubule and cortical collecting duct [CNT/CCD]) **[Figure 1b,c]**. There is no significant change in the percentage of proliferating (PCNA(+)) cells between vehicle- and furosemide-treated mice in PT **[Figure 1d]**. In TAL cells, there is very low PCNA detection compared to other cell types. In contrast, in the distal nephron, we detect statistically significant 4.0x-fold (4.08 ± 0.96, p-value=0.0169) and 5.5x-fold (5.45± 1.06, p-value=0.0033) increases in PCNA(+) cells in DCT and CNT/CCD principal cells, respectively, after one week of furosemide treatment. After three weeks of treatment, distal tubular cell hyperplasia is similar to vehicle-treated mice. Taken together, our results demonstrate that furosemide transiently induces cellular proliferation only in distal segments of the nephron. Interestingly, TAL cells exhibit minimal turnover compared with other cell types.

### Chronic furosemide treatment induces progressive proximal and distal hypertrophy along the nephron

After one week treatment, furosemide induced a modest, yet statistically significant increase in cell size of 3.8% (± 0.7, p-value=0.0031) in PT, 1.9% (± 0.6, p-value≤0.0001) in DCT, and 16.9% (± 0.7, p-value≤0.0001) in CNT/CCD principal cells [**Figure 2b, d**]. Furosemide inhibits NKCC2 in TAL. Therefore, we hypothesized that furosemide would decrease the size of cells in this segment. However, we detect no significant change in cell area of the TAL, -2.2% (± 0.8, p-value=0.1710) between treatment groups.

**Figure 2:**
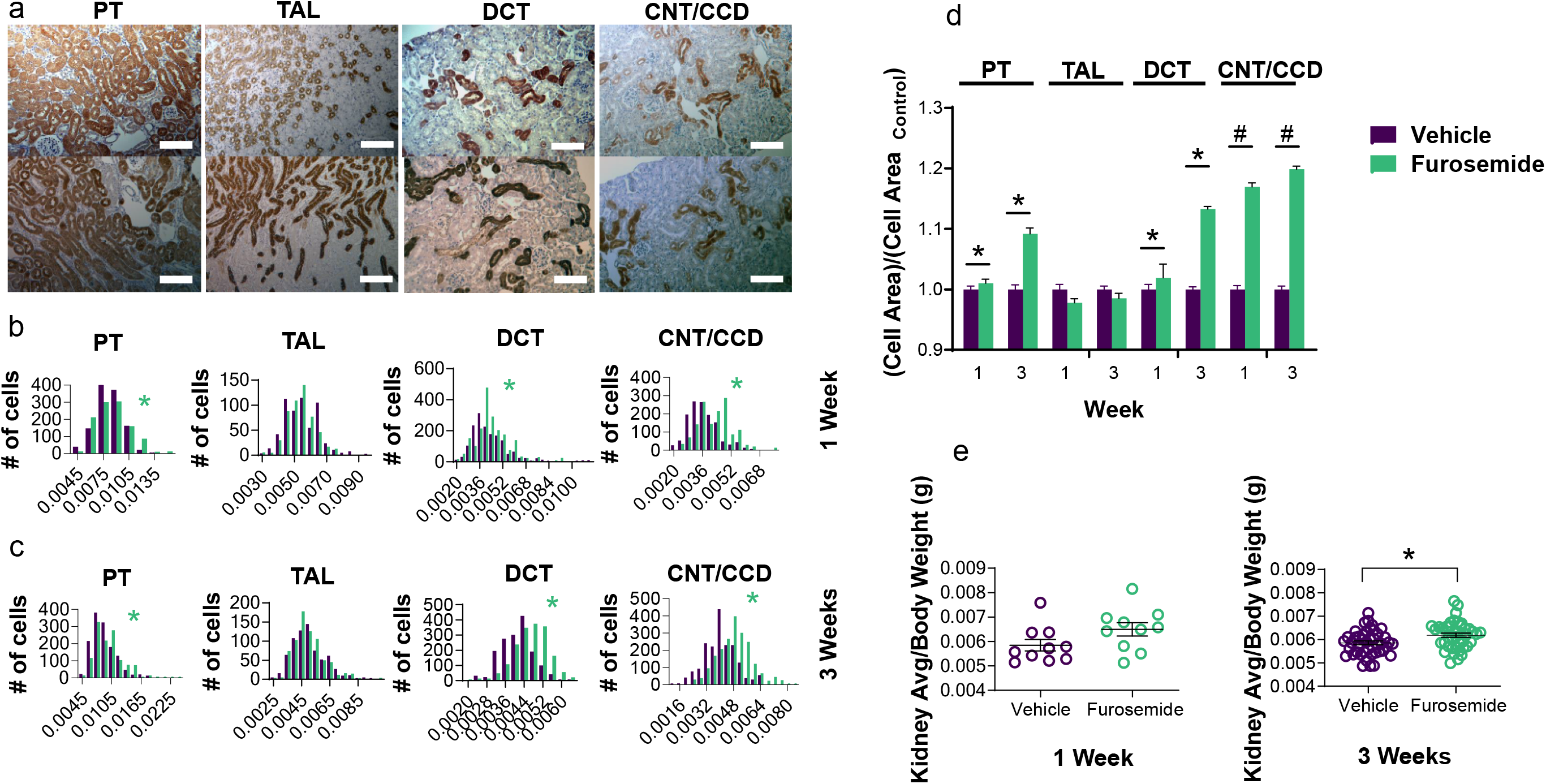
Furosemide induces progressive proximal and distal tubular hypertrophy. (a) Sample images of immunohistochemically stained vehicle controls (above) and furosemide treatment (below) using segment-specific markers. Scale bars: 100 µm. (b) PT, TAL, DCT, and CNT/CCD frequency histograms of weighted cell area values for mice fed vehicle or furosemide for one week. (N=5 mice per group, 150 tubules/group, n=531-2035 cell area values). Units are 300 pixels per inch. (c) PT, TAL, DCT, and CNT/CCD histograms of weighted values for mice treated for 3 weeks. (N=7 mice/group, 210 tubules/group; n=712-2045 cell area values). (d) Values were normalized to corresponding tubule/timeframe vehicle-treated groups. Histogram shows vehicle-(*purple*) and furosemide-treated (*green*) normalized values for PT, TAL, DCT, and CNT/CCD segments at 1 and 3 week time points (mean + SEM). (e) Global analysis of kidney growth was determined by the ratio of average kidney to body weight for one week-treated (*left panel*, N=10 mice/group) and three week-treated mice (*right panel*, N=41-43 mice/group). Data are presented as mean ± SEM. *p-value < 0.05 vs. vehicle treatment. #p-value < 0.05 vs. other nephron segments.

In contrast to the transient hyperplasia, three weeks of furosemide treatment resulted in progressively increased cellular hypertrophy in proximal and distal segments; PT, DCT, and CNT/CCD cell area is higher by 9.2%, 13.3%, and 19.9%, respectively [**Figure 2c, d**]. Moreover, furosemide increases the kidney-to-body weight ratio as early as after one week of treatment, and the difference between groups is significant at three weeks (5.85 ± 0.08 KW/BW vs. 6.16 ± 0.09 KW/BW, p-value=0.0084) **[Figure 2e]**. Similar to results with shorter-term treatment, we found no effect of three weeks of furosemide treatment on cell size in the TAL, - 1.4% (± 0.9, p-value=0.2332) **[Figure 2b-d]**. We also developed an orthogonal lectin-based flow cytometry method to examine hypertrophic changes at the single cell level **[Figure S2]**. Using this approach, we quantified a similar furosemide-induced increase in cell size in LTL-bound PT and DBA-bound CNT/CCD principal cells (9.3% ± 3.3, p-value=0.0203 and 13.8% ± 3.9, p-value=0.0071, respectively).

Taken together, these results establish that furosemide progressively increases the size of cells in proximal and distal segments of the nephron and increases kidney weight without reducing cell size in TAL.

### Segment-specific analysis using single cell RNAseq

We treated mice for one week with vehicle or furosemide, i.e., when we first detected both hyperplasia and hypertrophy. After optimizing for viable single cell isolation [**Figure S3**], we performed scRNA-seq of 24,711 cells from six mice (N=3 mice/group; **Table 1**). After accounting for error correction, removal of doublets, we normalized the number of cells across samples and analyzed the number of molecules per cell for 451 of the 500 genes across 22,154 cells.

**Table 1:** Quality control analysis of single cell RNAseq data. Vehicle-(*purple*) and furosemide-treated (*green*) mice. N=3 mice/group.

|  | Master Data Table | Mouse #1 | Mouse #2 | Mouse #3 | Mouse #4 | Mouse #5 | Mouse #6 | Pass Genes | Doublets Removed | Mixed Removed | Equal # Cells |
| --- | --- | --- | --- | --- | --- | --- | --- | --- | --- | --- | --- |
| Number of cells | 24711 | 4254 | 3628 | 3454 | 5281 | 3840 | 4254 | 24711 | 24282 | 23705 | 22154 |
| Number of genes in file |  | 500 | 500 | 500 | 500 | 500 | 500 |  |  |  |  |
| Number of genes in concatenated table | 500 | 500 | 500 | 500 | 500 | 500 | 500 | 456 | 456 | 456 | 456 |
| Number of genes detected | 491 | 479 | 478 | 478 | 477 | 476 | 465 | 451 | 451 | 451 | 451 |
| Mean molecules / cell | 2111 | 2129 | 2217 | 2332 | 2233 | 1805 | 1950 | 1630 | 1618 | 1584 | 1581 |
| Median molecules / cell | 1771 | 1794 | 1928 | 1913 | 1884 | 1454 | 1635 | 1336 | 1326 | 1304 | 1303 |
| Max molecules / cell | 13511 | 11253 | 11367 | 12712 | 13511 | 9907 | 10245 | 11485 | 11485 | 11485 | 11485 |
| Min molecules / cell | 182 | 296 | 417 | 483 | 429 | 346 | 182 | 12 | 12 | 12 | 12 |
| Mean genes / cell | 111 | 112 | 111 | 119 | 116 | 105 | 105 | 109 | 109 | 108 | 108 |
| Median genes/ cell | 110 | 111 | 110 | 119 | 114 | 103 | 102 | 108 | 108 | 107 | 107 |
| Max genes / cell | 224 | 220 | 213 | 215 | 224 | 223 | 208 | 221 | 221 | 221 | 221 |
| Min genes/ cell | 13 | 32 | 13 | 47 | 51 | 43 | 33 | 11 | 11 | 11 | 11 |

With visualization of single cell profiles in two dimensions using Uniform Manifold Approximation and Projection (UMAP) and based on known localization of select genes in the nephron^31-36^, we identify 15 kidney cell types for each of these clusters. The proximal tubule subsegments S1 through S3 segregate into two main clusters that contain overlapping and non-overlapping genes, we term S1/S2, S1/S2/S3, or S2/S3. While the descending limb of the loop of Henle segregates from proximal tubular cells, the thin and thick ascending limbs cluster together (TAL). DCT segregates into DCT1 and DCT2. We designed our single cell isolation protocol for tubular epithelial cells and utilized a 40 µm filter to eliminate cell clumps and glomeruli. However, we retain a small cell cluster with glomerular-specific genes. Although the thick ascending limb and CD are functionally segregated into cortical and medullary compartments^37^, initial analysis of TAL reveals one primary cluster relative to other cell types. On the other hand, the CD segregates into distinct clusters that we determined to be cortical principal cells (CCD), α-ICs, β-ICs, and medullary principal cells (MCD). We combined genes marked for lymphocytes and macrophages into an immune cell category. In [**Figures 3b,3c**], representative genes to identify each cell type are shown. After cell type identification, we compare the projections of cells from vehicle-vs. furosemide-treated mice. Analysis of cell proportions indicate no obvious batch effects related to yield of different nephron cell clusters [**Figure S4**] within or across samples.

**Figure 3:**
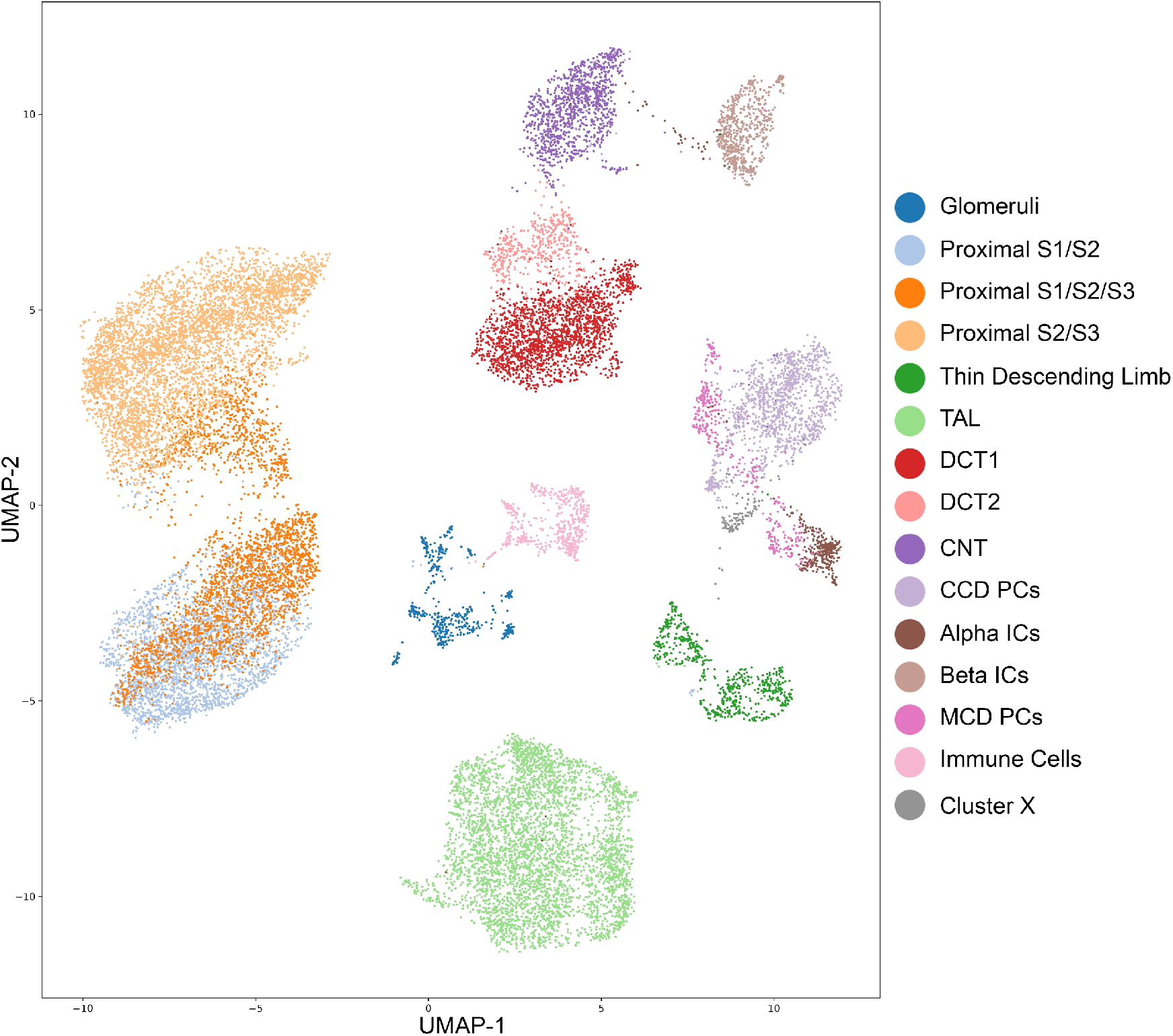
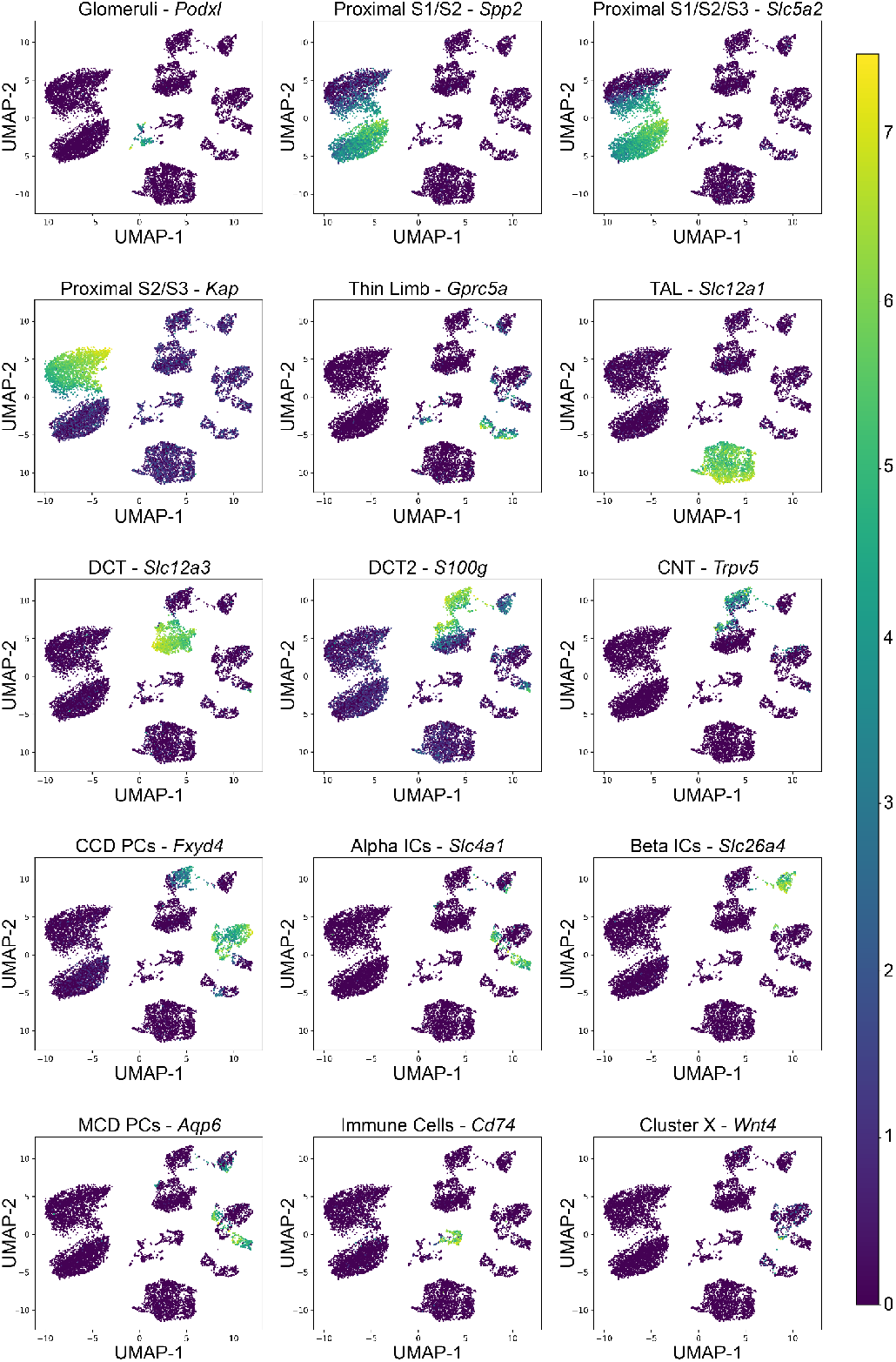
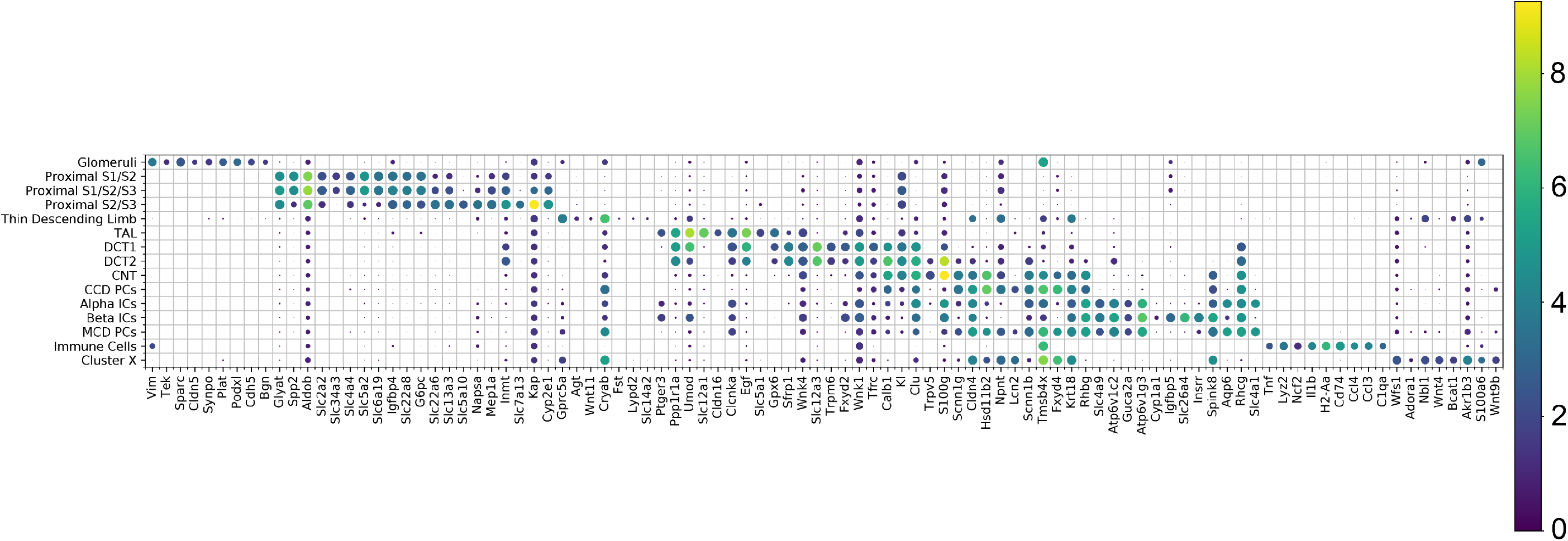
Unbiased clustering of nephron cell types in vehicle- and furosemide-treated mice. (a) Using the UMAP algorithm, gene expression across 451 target genes from 22,154 cells clustered into 15 subtypes that corresponded to known nephron segments and cell types and one additional cluster, we termed ‘Cluster X’. (b) For each of 15 clusters the most abundant genes and their intensity and distribution of expression across all cell types on a UMAP projection. This expression pattern is an aggregate of six total vehicle- and furosemide-treated mice (N=3 mice/group). (c) Nephron-wide distribution of expression of most abundant genes within each cluster. The scale for molecules per cell is a logarithm scale ranging from highest molecules/cell (*yellow*) to lowest (*purple*).

### Furosemide reduces TAL and increases DCT cells

While the majority of cell types are similar in cell number between vehicle- and furosemide-treated mice, we detect significant changes between several cell clusters [**Table S2**]. Two groups, the TAL and DCT2, have the largest fold decrease and increase, respectively. The subpopulation of TAL cells in furosemide-treated mice are 43% less than the identified TAL subpopulation in vehicle-treated mice (874.7 ± 43.4 vs. 499.3 ± 101.7, p-value=0.001, FDR=0.017). Congruent with the hyperplasia in morphometry experiments, cells identified as DCT2 increase ∼2.0-fold in cell number in furosemide-treated mice (53.3 ± 10.9 vs. 108.3 ± 2.0, p-value=0.012, FDR=0.082). We also observe similar differences when we measure by the proportion of total cells between vehicle- and furosemide-treated mice [**Figure 4, Figure S5, Table S3**].

**Figure 4:**
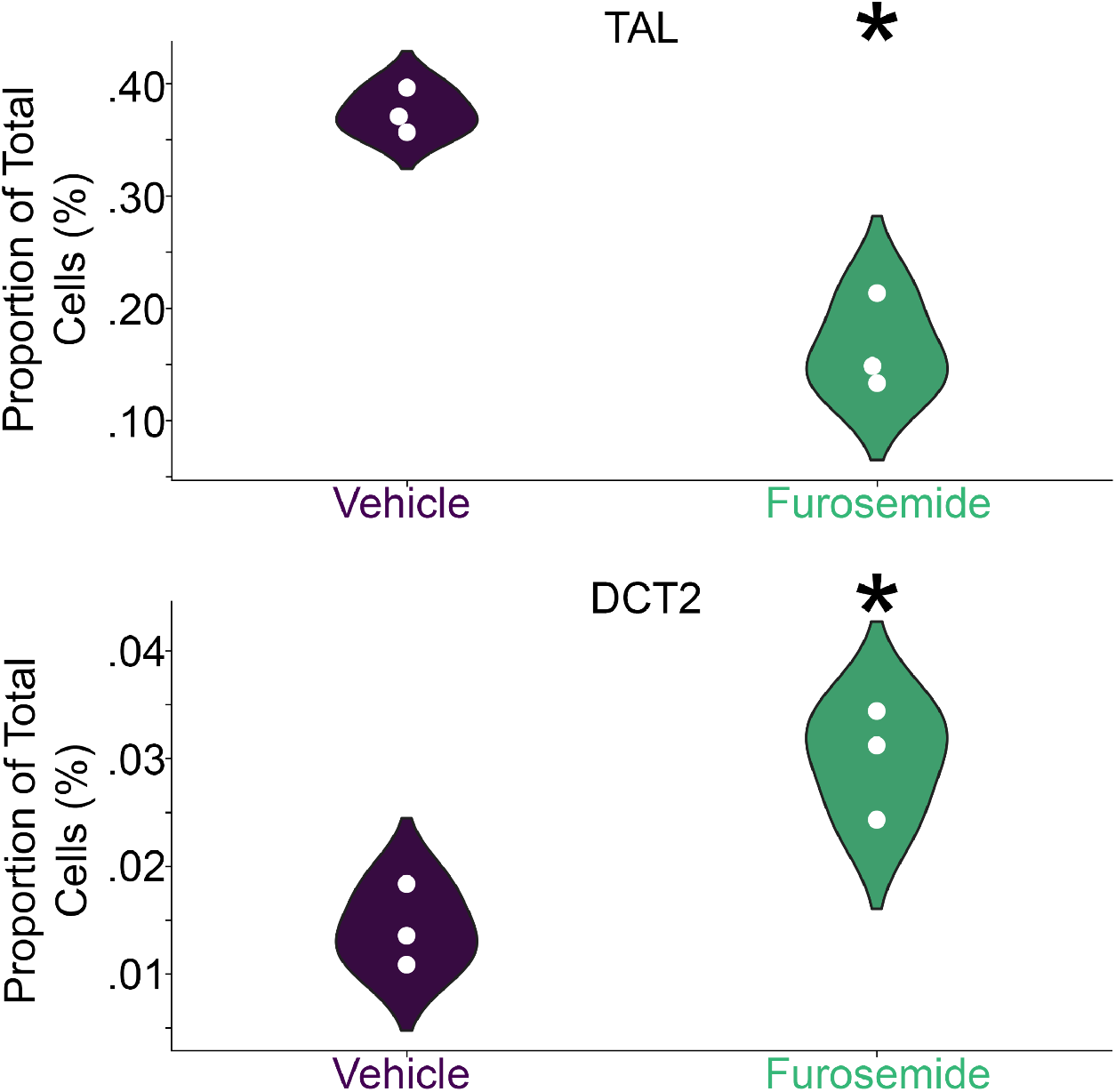
Smaller and larger proportions of distinct cell populations with one week furosemide treatment. Proportion of cells per cluster (segment/cell type) in vehicle-(*purple*) and furosemide-treated (*green*) mice. Each dot represents one mouse. Data is represented by violin plots. *, indicates cell clusters with false discovery rate < 0.1. TAL, thick ascending limb; DCT2, distal convoluted tubule segment 2. N=3 mice/group.

### Differential gene expression across cell types and phenotypes

We identify 287 genes that are significantly up- or down-regulated within a specific cluster, and changes in gene expression are distinct when compared across cell clusters. A heat map of genes with the largest effect sizes between vehicle- and furosemide-treated mice and the relative expression along the nephron is shown in [**Figure 5a**]. The number of genes that are significantly different within a cell cluster vary along the nephron. As shown in [**Figure 5b**], the proximal tubule and the TAL demonstrate the largest number of differentially regulated genes and the largest cell clusters. The cell clusters with the largest change relative to the number of cells are the thin limb, MCD PCs, and Cluster X, which we termed *Wnt4*(+) cells based on the specific pattern of gene expression. Cell clusters with least the change relative to the population of each cluster are proximal tubular cell types, S1, S1/S2/S3, and S2/S3, and alpha intercalated cells. Several genes are up- or down-regulated in multiple nephron cell types. For example, *Tfrc*, which encodes for the transferrin receptor 1^38^, is significantly up-regulated in nine cell clusters and down-regulated exclusively in TAL, while *Smad7*^39^, is down-regulated in eight cell clusters. The alpha subunit of the Na-K ATPase, *Atp1a1*, is also up-regulated in several sodium-transporting nephron segments. A rank order of genes that are altered across cell types is shown in [**Figure 5c**]. The genes with the largest effect sizes in any cell cluster are shown in [**Figure 5d**]. Compared to modest changes in genes that are altered in multiple cell clusters, genes with large effect sizes are mostly altered in one or only a few cell clusters. For example, *Birc5*, a hyperplasia gene is increased in a small subset of DCT1 cells (z-score=2.703, FDR=0.0052), and *Ptgs2*, a cyclo-oxygenase involved in NKCC2 inhibition and natriuresis^40^, is decreased primarily in CCD principal cells (z-score=-1.659, FDR=0.0004).

**Figure 5:**
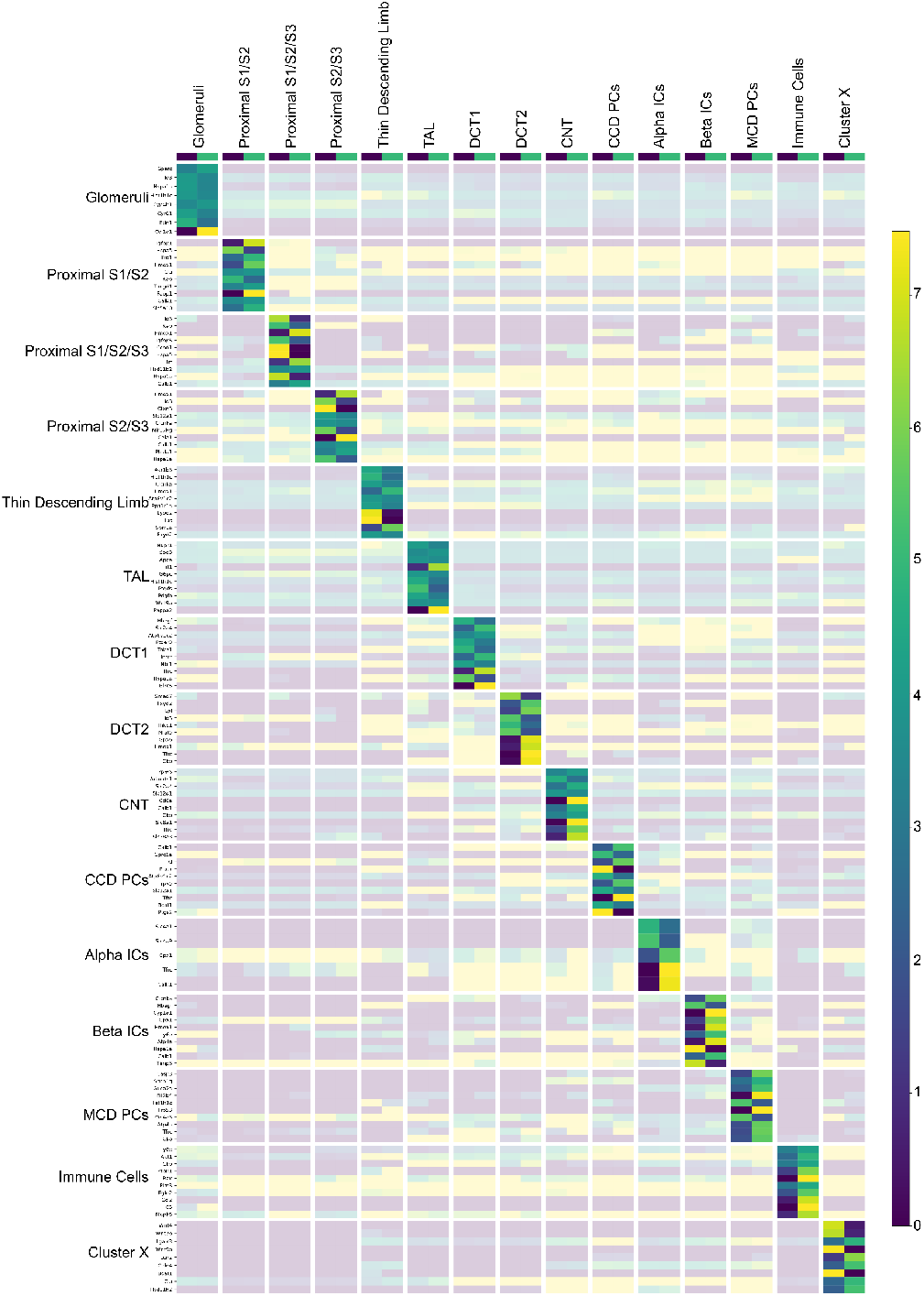
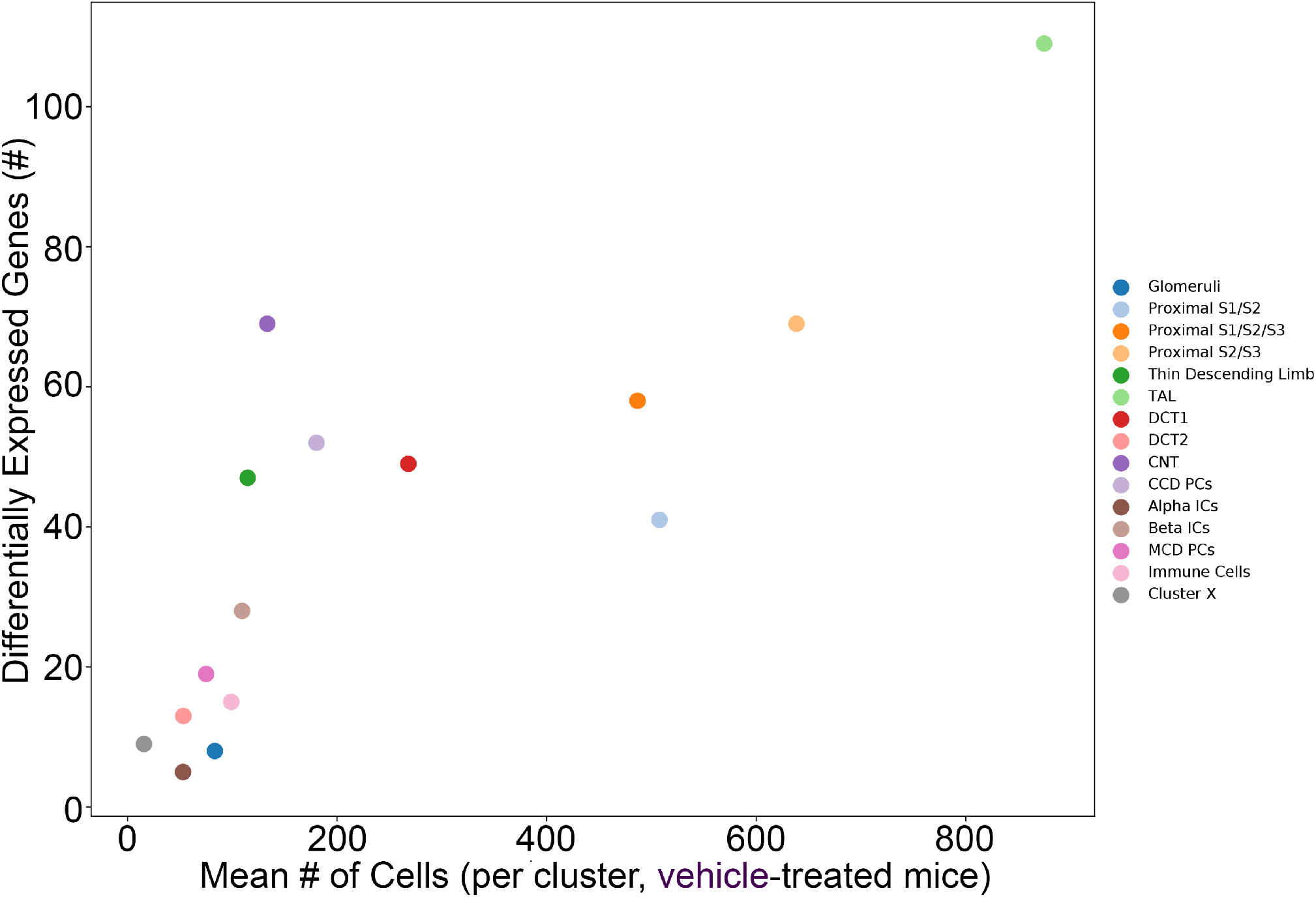
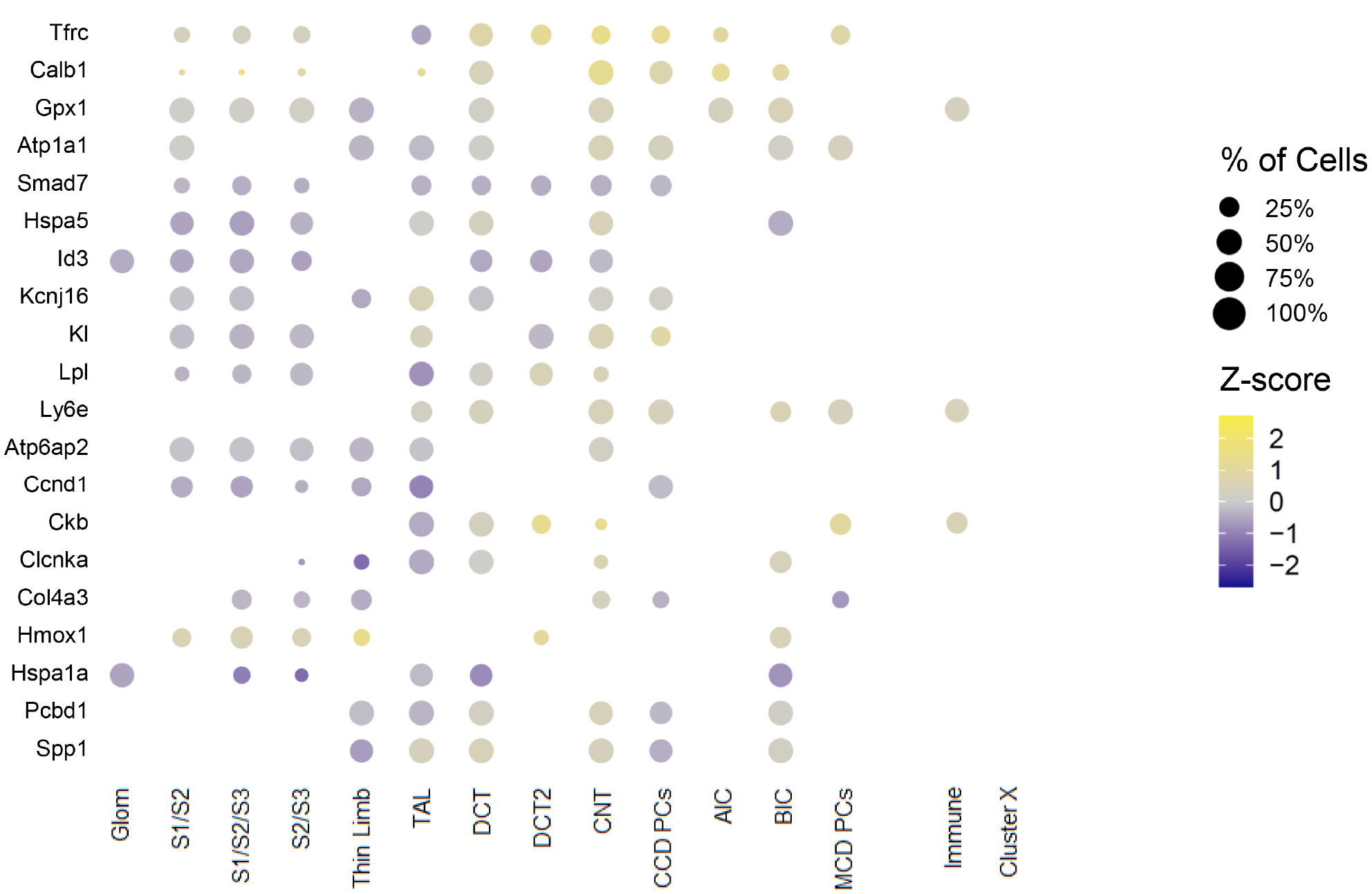
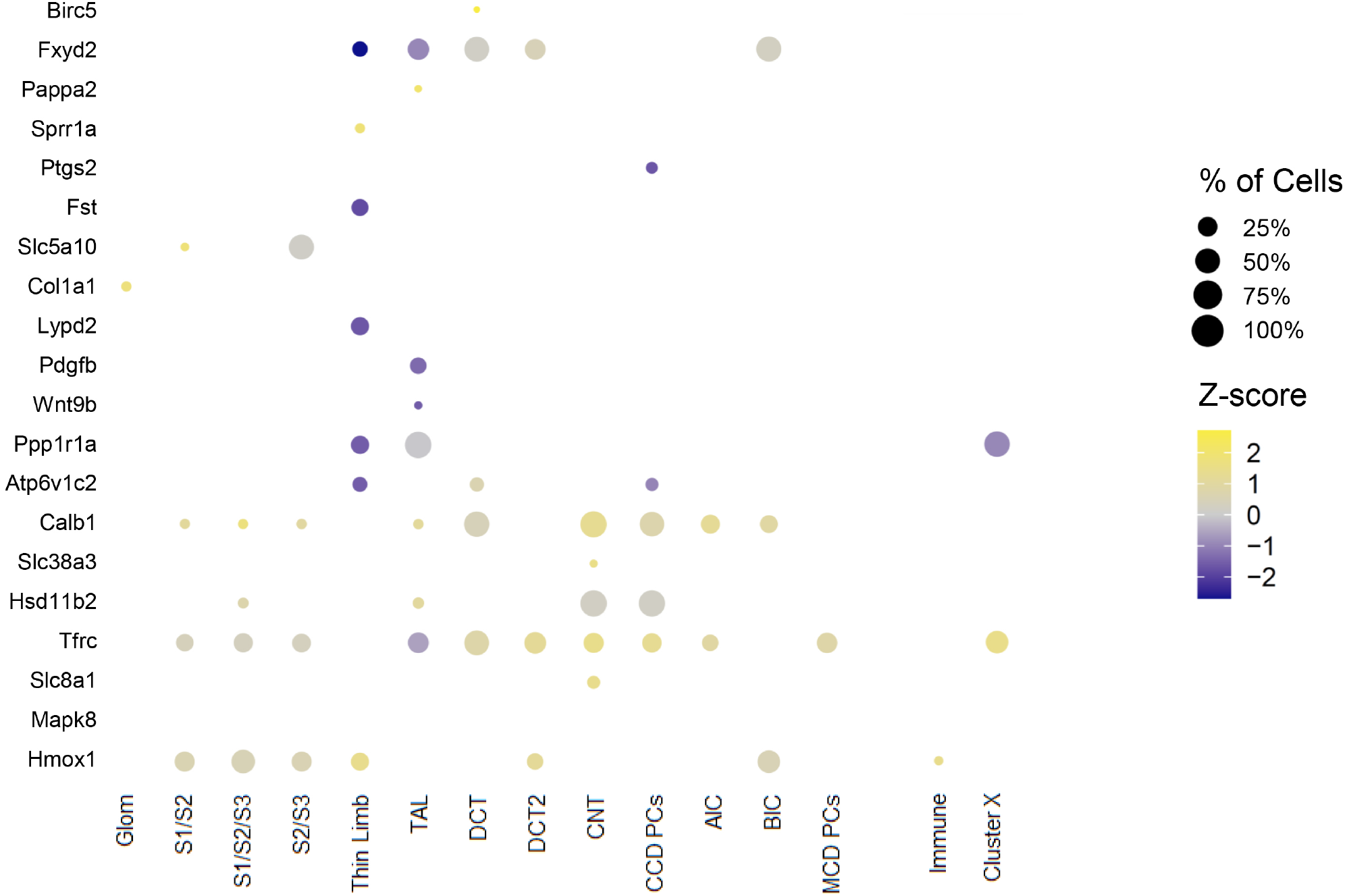
Differentially expressed genes within cell clusters. (a) The differentially expressed genes between cells from vehicle- and furosemide-treated mice, with a low false discovery rate and the highest absolute effect sizes are shown on the left for each segment. The relative expression level of each of these genes is shown across all cells and arranged by treatment-type (*legend*, vehicle [*purple*] and furosemide [*green*]) and cell cluster (*top, left*) as shown. Data are presented as a heat map (*scale, right*) by mean expression per cluster from N=22,154 cells clustered into 15 subtypes from N=3 mice/group. (b) To assess for enrichment of differential expression within a cell cluster, we graphed the number of differentially expressed genes per mean population size of each cluster (in vehicle-treated mice). (c) Differentially expressed genes ranked by highest frequency of cell clusters. The color of the dot refers to the effect size (see heat map, *right*), and the size of the dot is proportional to the number of cells expressing that gene in a given cell cluster (see scale, *right*). (d) Differentially expressed genes ranked by highest effect size within a cell cluster. The color of the dot refers to the effect size (see heat map, *right*), and the size of the dot is proportional to the number of cells expressing that gene in a given segment (see scale, *right*).

### Furosemide increases distal nephron-specific ion transport along the nephron

As depicted in [**Figure 6, Table S1**], several of the differentially expressed genes in the distal nephron related to sodium transport pathways are altered in each of these nephron segments. The directionality of the changes is congruent with the known compensatory increase in sodium transport activity in distal (vs. proximal) nephron segments with furosemide therapy^12,13,15,41^. The genes that are up-regulated differ in each cluster, and along the axis of the distal nephron from the TAL to DCT1 to CCD PCs, there is a gradual increase in the up-regulation of genes related to increased sodium reabsorption. From DCT2 onward, the only down-regulated gene in these pathways is *Nedd4-2*, which represents disinhibition of NCC-^42,43^ and ENaC-mediated^44-47^ sodium transport. In contrast, *Slc9a3, Slc5a2, Slc34a2*, and *Slc34a3*, genes that encode for major proximal tubular sodium transport proteins (NHE3, SGLT2, and NaPi-2, respectively) are not up-regulated in their respective segments [**Table S1**].

**Figure 6:**
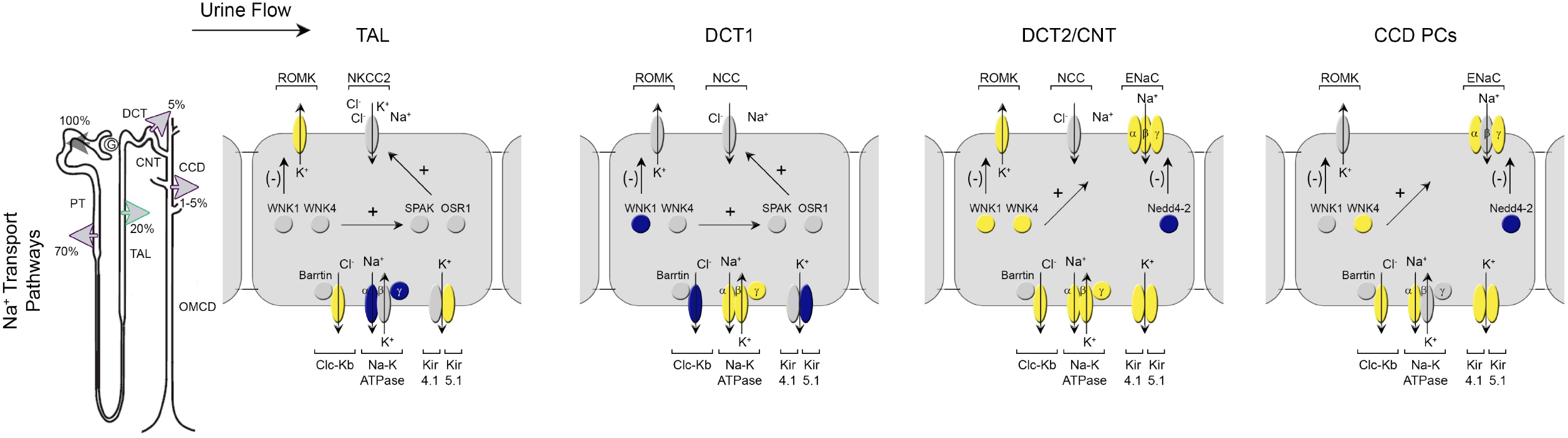
Differentially expressed genes related to mechanisms of ion transport in the distal nephron. Differentially expressed genes that are significantly different within a given cell cluster from vehicle-vs. furosemide-treated mice are represented as up-(*yellow*) or down (*blue*)-regulated (*right*). Genes known to be part of sodium transport within these cells but are not differentially regulated are showed as neutral (*gray*). *Left panel*, the proportional amount of solute reabsorption is shown along the nephron. Genes that are not detected or represented by scRNAseq are shown as transparent. Each cell cluster is represented by successive panels left to right: TAL, DCT1, DCT2/CNT, and CCD PCs as indicated. Due to similarities in function and expression, DCT2 and CNT cells are represented by a single diagram. TAL, thick ascending limb, DCT1, distal convoluted tubule segment 1, DCT2, distal convoluted tubule segment 2, CNT, connecting tubule, CCD PCs, principal cells of the cortical collecting duct.

### Furosemide increases IGF-1R-dependent kidney weight and proximal, but not distal, tubular cell hypertrophy

To better understand mechanisms of tubular remodeling, we focused on hypertrophy. Furosemide induces IGF-1 protein expression in rat kidney^24^, but its contribution to cellular hypertrophy is unknown. Consistent with earlier studies, whole kidney samples from furosemide-treated mice demonstrated a 7.4-± 1.3-fold (p-value= 0.0029) increase in IGF-1 protein band intensity relative to vehicle [**Figure S6**]. Also, the expression of *Pappa2*, that encodes a secreted IGF binding protein protease that increases IGF-1 bioavailability^48^, has one of the largest effect sizes with furosemide (z-score=2.054, FDR=9*10^−9^). These data suggest a higher propensity of IGF-1 receptor (IGF-1R)-mediated signaling with furosemide treatment. Thus, we generated a novel, inducible renal tubule IGF-1R knockout mouse model (KO). As expected from the breeding scheme, approximately one-quarter of pups were *Igf1r*^Pax8/TetOCre^ (KO) mice. Depicted in **Figure 7A**, *Igf1r*^Pax8/TetOCre^ mice were homozygous for the floxed IGF-1 receptor allele and hemizygous for both the Pax8-rtTA and TetOCre transgenes. We deleted IGF-1R in fully mature 12-week old mice to avoid confounding of impaired IGF-1-mediated renal development on the kidney-to-body weight ratio^49,50^. After two weeks of doxycycline, immunoblots demonstrate a marked reduction in IGF-1 receptor abundance in whole kidney lysates of KO compared to littermate controls (control 1.00 ± 0.14 vs. KO 0.11 ± 0.04 arbitrary units, p-value< 0.0001) [**Figure 7B**]. We measured body weight between KO and littermate controls after doxycycline treatment and there is no divergence in their growth curves [**Figure 7C**].

**Figure 7:**
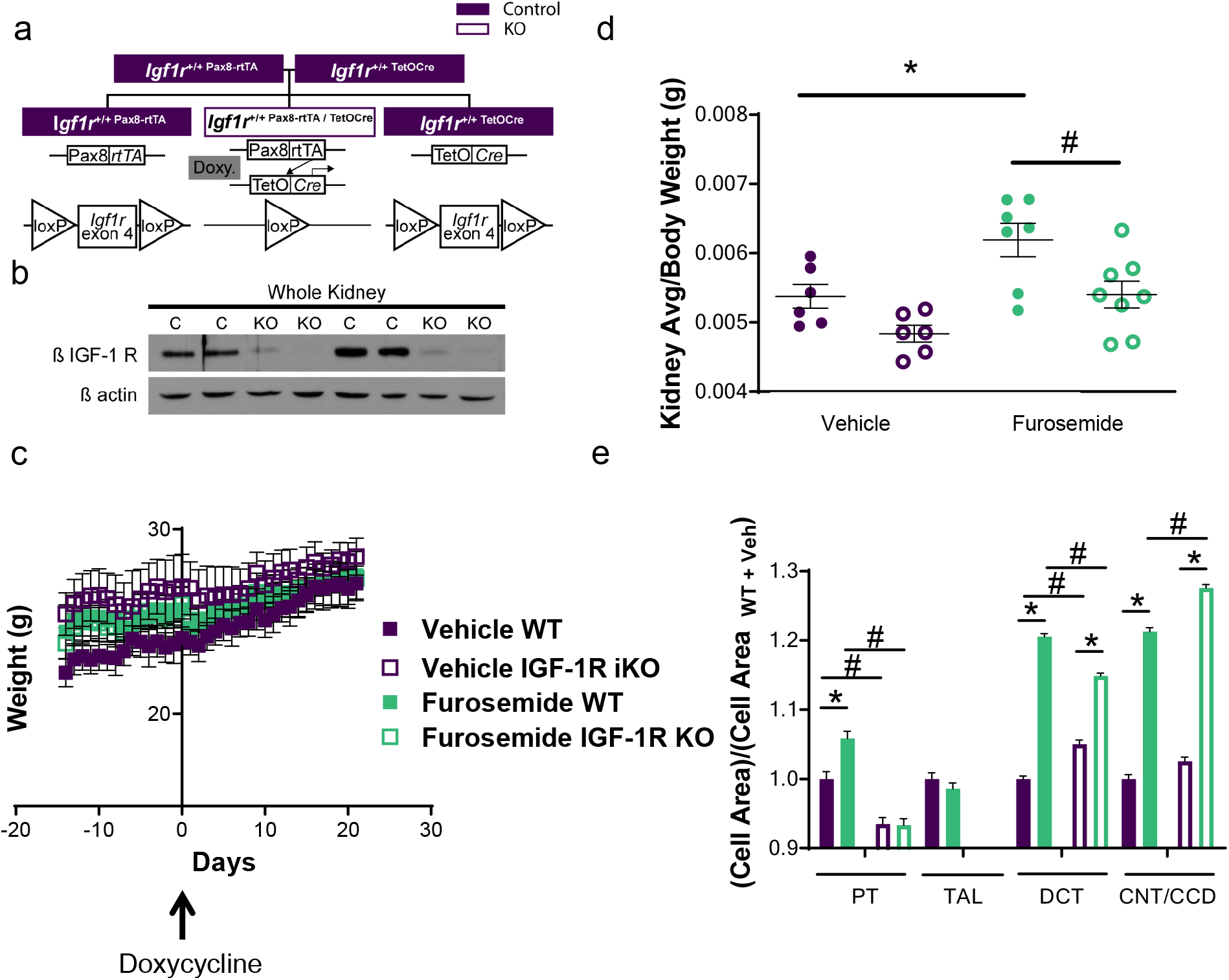
Inducible deletion of renal tubular Insulin-like growth factor 1 (IGF-1) receptor reduces furosemide-induced kidney growth and proximal tubular hypertrophy. (a) Breeding strategy and schematic of tetracycline-inducible Cre-loxP system to generate inducible renal tubular IGF-1 receptor knockout (KO) mice. (b) Representative immunoblot of whole kidney lysates probed for the ß subunit of the IGF-1 receptor (ß IGF-1R) four weeks after completing two weeks of doxycycline administration, then stripped and reprobed for ß actin. (c) Growth curves for control and KO mice before and after doxycycline administration. (d) Kidney-to-body weight ratio of vehicle- and furosemide-treated control and KO mice (N=6-8 mice/group). (e) Cell areas from three-week vehicle- and furosemide-treated control and KO mice. Values were normalized to corresponding tubule vehicle-treated controls. (N=7 mice/group, 210 tubules/group; n=531-2045 cell area values). Control, age and sex-matched littermates of KO mice; Doxy., doxycycline. Data are presented as mean ± SEM. *p-value < 0.05 vs. vehicle treatment; #p-value < 0.05 vs. littermate controls.

Deletion of IGF-1R in fully mature renal tubular epithelium prevents the increase in kidney-to-body weight of furosemide-treated mice [**Figure 7D**]. There is no significant difference in PT cell area between vehicle-vs. furosemide-treated KO mice. However, these KO mice still develop significant hypertrophy in DCT and CNT/CCD principal cells comparable or higher to littermate controls [**Figure 7E**]. These data suggest that furosemide increases kidney weight and selectively induces tubular hypertrophy in the proximal tubule via an IGF-1R-mediated pathway. In contrast, furosemide-induced distal tubular hypertrophy is independent of the IGF-1R.

### Proximal tubule changes in genes upstream and distal tubule changes downstream IGF-1R

We next analyzed for cell type-specific changes in hypertrophic pathways. As depicted in [**Figure 8**], in vehicle-vs. furosemide-treated mice, modulation of genes upstream of the IGF-1R, i.e. IGF binding proteins^51^ (*Igfbp1, Igfbp5*) are confined to proximal tubular cell types, S1-S3. In contrast, genes downstream of IGF-1R, e.g. PI3-kinase pathway genes (*Pdk1*^*52*^, *Akt1*^53-55^, *FKBP4*^*56*^, *Eif2BP4*^*57,58*^) are up-regulated in distal nephron segments (CNT, CCD) and *Foxo3*, an inhibitory transcription factor downstream of PI3-kinase^*59*^, is down-regulated in TAL and DCT1. *Spp1*, which encodes for osteopontin, a pro-hypertrophic secreted cytokine that acts, in part, by promoting PI3-kinase^60-62^, is also up-regulated in selected distal nephron segments (DCT, CNT). In contrast, PI3-kinase pathway genes are not up-regulated in proximal tubular cells despite significant hypertrophy.

**Figure 8:**
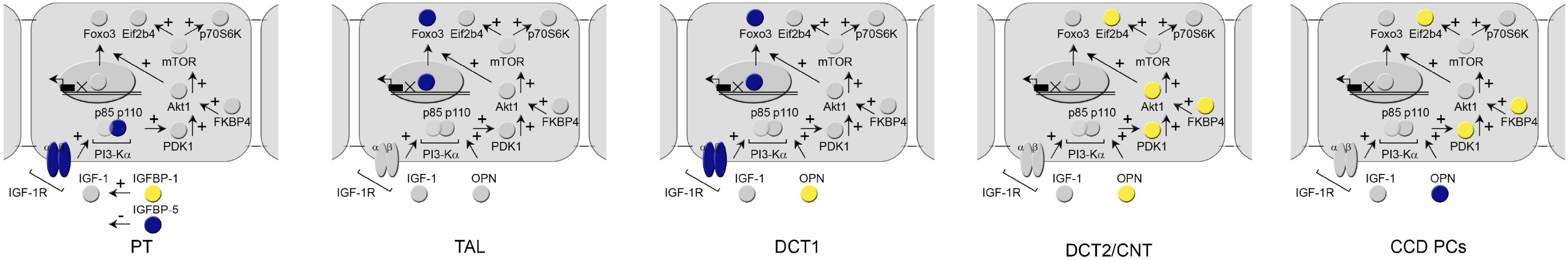
Differentially expressed genes related to mechanisms of hypertrophy along the nephron. Differentially expressed genes that are significantly different within a given cell cluster from vehicle-vs. furosemide-treated mice are represented as up-(*yellow*) or down (*blue*)-regulated (*right*). Genes known to be part of PI3-kinase-mediated hypertrophy pathways within these cells but are not differentially regulated are showed as neutral (*gray*). *Left panel*, the proportional amount of solute reabsorption is shown along the nephron. Genes that are not detected or represented by scRNAseq are shown as transparent. Each cell cluster is represented by successive panels left to right: PT, TAL, DCT1, DCT2/CNT, and CCD PCs as indicated. Due to similarities in function and expression, S1, S2, and S3 are represented by a single diagram and DCT2 and CNT cells are represented by a single diagram. PT, proximal tubule TAL, thin and thick ascending limb, DCT1, distal convoluted tubule segment 1, DCT2, distal convoluted tubule segment 2, CNT, connecting tubule, CCD PCs, principal cells of the cortical collecting duct.

## DISCUSSION

### Nephron plasticity is common and consequential yet poorly understood

Furosemide therapy is commonly prescribed. Twenty million prescriptions for furosemide were tabulated in 2015 alone^63^, and furosemide is on the World Health Organization list of essential medicines for provision of health care^64^. In this series of studies, we use a common therapeutic– furosemide– to examine consequences to specific cell types along the entire nephron using single cell analysis. These effects have relevance to the care of patients who receive loop diuretic therapy, including those with heart failure, chronic kidney disease, and/or other edematous states. Remodeling of the proximal or distal tubule as observed with furosemide also occurs in disease states such as diabetes, unilateral nephrectomy, and hypokalemia^65-67^. However, the mechanisms of this functionally and clinically relevant nephron plasticity are unknown.

### Proximal vs. Distal Tubular Remodeling

Using morphometry, flow cytometry, inducible cell-specific gene deletion, and single cell RNAseq, we found that furosemide progressively increases hypertrophy in the proximal tubule and kidney size that is dependent on tubular epithelial IGF-1R. The proximal tubule undergoes hypertrophy without significant hyperplasia, similar to angiotensin II (Ang II) induced hypertrophy of proximal tubular cells^68,69^ or compensatory renal growth associated with uninephrectomized mice or rats^70^. The full functional consequences of proximal tubular hypertrophy to solute transport remain to be explored. Furosemide induces renin release but diminishes proximal tubular sodium reabsorption^71,72^ in the setting of sodium repletion. It is noteworthy that we detect proximal tubular cell hypertrophy under similar conditions of sodium repletion and either little to no change (e.g., *Slc9a3, Slc5a2, Slc34a2*, and *Slc34a3*) or down-regulation of sodium transport machinery (e.g. subunits of the Na-K ATPase pump, *Atp1b1*). These results suggest that tubular hypertrophy may be, in part, independent of enhanced sodium reabsorption^73^ and would further differentiate the mechanisms of hypertrophy in different nephron segments.

In contrast to proximal tubular cells, distal tubular cells comprise only a small percentage of nephron mass, but exhibit both hyperplasia and progressive hypertrophy, independent of IGF-1R. Distal tubular remodeling is already known to have functional consequences for diuretic resistance. Individuals treated with furosemide have significantly increased sensitivity to thiazide and thiazide-type diuretics^12^, providing the rationale for dual class diuretic therapy in states of refractory edema^14,74^. Distal tubular hyperplasia is supported by morphometry for proliferating cells, and a significantly increased number of cells by single cell RNAseq, with higher expression of hyperplasia-specific genes, e.g. *Birc5*, an anti-apoptotic gene and pro-growth gene^75^. Although there are clinical studies measuring increased ^76^ or decreased^77^ proliferation of cancer cells due to furosemide treatment, to our knowledge this is the first report of distal tubular hyperplasia. Furthermore, distal tubular segments are enriched for changes in ion transport genes with furosemide treatment. Our data show similarities and differences in mechanisms of diuretic resistance. The degree of hyperplasia and hypertrophy differ in DCT and CNT/CCD PCs, and transcriptional changes in the same genes, albeit congruent with enhanced distal sodium transport, differed by nephron cell type from the thick ascending limb to the cortical collecting duct. Taken together, these data are consistent with human physiology studies that the compensatory increase in sodium transport with furosemide is primarily in distal vs. proximal tubule^13^ and highlight precise nephron-segment specific mechanisms.

### Changes in cell abundance with diuretic therapy

The TAL show no hyperplasia or changes in cell size (atrophy or hypertrophy) at one or three weeks of treatment; however, this segment had the highest number of differentially expressed genes with furosemide treatment. We also detect significantly decreased numbers of TAL cells and increased numbers of DCT2 cells by single cell RNAseq. These furosemide-induced changes could be the result of transdifferentiation from one cell type to another, or simply an artifact of the cell dissociation process. However, three independent pieces of evidence suggest that indeed, TAL cells decrease and DCT2 cells are more abundant with furosemide. One, DCT cells display more PCNA(+) nuclei with treatment, suggesting proliferation rather than conversion from TAL cells. Two, the number of TAL cells that decrease are fewer than the increased number of DCT1 and DCT2 cells. Three, we do not see tissue staining for DCT-specific markers in the thick limb (data not shown).

Following hyperplasia, we measure progressive hypertrophy. This hyperplasia-to-hypertrophy progression in tubular remodeling is similar to studies of diabetes-associated hypertrophy^78-80^. Enhanced thiazide sensitivity and increased NCC protein^12,81^ could be due primarily to stimulation of NCC or related gene transcription and/or increased distal tubular mass with normal transport activity proportional to cell number and volume. Our data provide strong support for a role of remodeling. Interestingly, we detect an increase in sodium transport machinery (*Atp1a1, Clcnkb, Kcnj10*), but we do not detect an increase in molecules per cell of NCC (*Slc12a3*) in either DCT1 or DCT2 cells [**Figure 6, Table S1**].

### IGF-1 signaling, hypertrophy, and kidney size

Elevated IGF-1 protein in kidneys undergoing hypertrophy due to a variety of stimuli, including furosemide^24,29,82,83^, has been described before. Pappa2, an IGF-1 binding protein protease, known to increase IGF-1 bioavailability is up-regulated in TAL cells. Also, furosemide treatment increases the kidney-to-body weight ratio of rats ^22,23,72^ and now, mice, and the proximal tubule comprises the bulk of nephron mass (by immunofluorescence and flow cytometry, data not shown). Thus, we postulated that a furosemide-induced increase in proximal tubule cell size would be necessary and sufficient to increase kidney size. Indeed, IGF-1R deletion significantly attenuates the kidney-to-body weight ratio and proximal tubular hypertrophy. By differential expression, IGF-1 pathway genes are significantly enriched in proximal vs. distal tubular cells, consistent with our 3-week morphometry data.

These data inform our understanding of tubular hypertrophy. Using proximal tubular specific *Egfr* and *Pten* knockout mice, Harris and colleagues showed that tonic PI3-kinase signaling can override anti-proliferative genes in unstimulated kidneys^84^. In their model, unilateral nephrectomy further increased contralateral kidney size implying that additional pathways can generate a hypertrophic response. Our data suggest that in the proximal tubule, IGF-1R signaling is a potential alternative pathway for hypertrophy. These data have implications for kidney repair in the setting of acute or chronic injury when tubular proliferation may be warranted^85,86^. Moreover, targeting the IGF-1R to minimize potentially unwanted hypertrophy in the setting of diuretic resistance or other hypertrophic stimuli^87^, would not limit distal tubular hypertrophy. Differential expression at one week demonstrate specificity for changes in genes for IGF binding proteins^51^, *Igfbp1* and *Igfbp5* in proximal, but not distal tubular cell types. While IGF binding proteins can have different roles in different contexts, IGFBP-1 is known to enhance IGF-1 signaling *in vivo*^24,88^. Alternatively, the full-length form of IGFBP-5 can reduce IGF-1 bioavailability, and therefore IGF-1R-mediated signaling^89-92^. In contrast to modulation of upstream genes, genes downstream of IGF-1R (*Pdk1*^*52*^, *Akt1*^53-55^, *FKBP4*^*56*^, *Eif2BP4*^*57,58*^, *Spp1*^60-62^, *Foxo3*)^59,93^ are congruent with enhanced PI3-kinase signaling in distal but not proximal tubular cell types. These data highlight potential growth pathways that bypass the IGF-1R and demonstrate the heterogeneity of hypertrophic mechanisms in diuretic resistance.

### Limitations

We study tubular remodeling in mice which may limit translation to human studies. However, patients on furosemide rarely undergo kidney biopsy or nephrectomy and therefore mouse models provide a vehicle for comprehensive study of mechanisms along the nephron. Moreover, temporal and selective deletion of IGF-1 signaling in the kidney is not feasible in patients treated with furosemide. Effects of furosemide may be subject to confounding by indication, and thus, we provide high dose diuretic therapy in the absence of edema-forming states such as heart failure, cirrhosis, or nephrotic syndrome. Further studies of furosemide therapy in these conditions will be needed to confirm our findings. For single cell RNAseq analysis we chose a targeted gene panel and thus, may have missed some differential gene expression. However, with one week of therapy and focused on 451 genes, we undercover a robust network of changes in gene expression in specific cell clusters and changes in cell proportions and elucidate multiple, novel mechanisms of tubular remodeling.

### Conclusions

Using single cell morphometry and RNA sequencing, we demonstrate cell type-specific forms of tubular remodeling including selective changes in hyperplasia, distinct IGF-1 receptor-dependent and independent hypertrophy, gain and loss of select cell types, and heterogeneity of differential gene expression across cell clusters. These findings highlight mechanisms of diuretic resistance, and have the potential to help scientists and clinicians develop new strategies to optimize care in edematous disorders.

## MATERIALS AND METHODS

### Study Approval

We performed all mouse experiments in accordance with the Guide for the Care and Use of Laboratory Animals (National Academies Press, 2011) and obtained approval from the Stanford University Institutional Animal Care and Use Committee.

### Vehicle and furosemide treatment

We fed mice a high-sodium (104 µmoles Na+ per kcal) gel diet with 70 mg/kg furosemide body weight or the volumetric equivalent of vehicle (ethanolamine) for one or three weeks before sacrifice as described previously^94^. We maintained control and mutant iT-IGF-1R KO C57BL/6J mice in individual cages for 1 – 3 weeks on a 12-h light/12-h dark cycle. We also monitored body weight and food intake daily. To detect the effects of furosemide, we compared daily water intake between groups for a single experiment [**Figure S1**] and every 2-3 days for all subsequent experiments.

### Antibodies and lectins

We utilized rabbit polyclonal antibodies for detection of endogenous IGF-1 (Santa Cruz Biotechnology, Santa Cruz, CA), the ß-subunit of the IGF-1 receptor (Proteintech Group, Rosemont, IL), Proliferation Cell Nuclear Antigen (PCNA) (Santa Cruz Biotechnology), Tamm-Horsfall protein (Santa Cruz Biotechnology); Parvalbumin (ThermoScientific, Waltham, MA); and Calbindin (EMD Millipore, Hayward, CA). We used a mouse monoclonal antibody to probe for ß-actin (EMD Millipore). We purchased all lectin conjugates from Vector Laboratories (Burlingame, CA).

### Tubular Morphometry Studies

We deparaffinized slides of 5 µm kidney sections in xylene, rehydrated in graded ethanol and washed in PBS buffer. For antigen retrieval, we boiled samples in 10 mM citrate buffer pH 6.0 for 10 minutes. We then blocked each sample with 5% v/v goat serum in suppressor serum (3% BSA in PBS-T) followed by overnight incubation of primary antibody at 4°C. We prevented nonspecific binding of biotinylated secondary antibodies with an Avidin/Biotin blocking kit (Vector Laboratories, Burlingame, CA). We incubated goat anti-rabbi HRP-conjugated secondary antibodies in a humidified chamber for 45 minute at 1:200 dilution and quenched endogenous peroxidase activity by a 20 minute incubation of 3% H_2_O_2_. We administered a Vectastain ABC complex kit (Vector Laboratories), followed by staining with 0.05% diaminobenzidine (DAB) in the presence of 0.015% H_2_O_2_ in PBS [pH 7.2]. To visualize nuclei, we stained slides with Mayer’s hematoxylin. In parallel, we performed immunohistochemical staining of samples without primary antibody to check for nonspecific binding of secondary antibodies. We captured images using a Leica light microscope and arranged them using Adobe Photoshop (Adobe Systems, San Jose, CA).

For immunofluorescence, we blocked slides with 10% normal donkey serum in suppressor serum before overnight incubation of primary antibodies or lectins. We again used Avidin/Biotin blocking kit (Vector Laboratories, Burlingame, CA) to prevent nonspecific binding of biotinylated lectins or streptavidin secondary antibodies. We incubated secondary antibodies for 45 minutes in the dark inside a humidified chamber. To visualize nuclei, we washed samples and used Vectashield Hard Set Mounting Medium with DAPI (Vector Laboratories).

To measure hyperplasia, we used immunofluorescence to detect PCNA in kidney samples^95,96^. To detect proximal tubule, thick ascending limb, DCT1, and principal cells of the CNT/CCD, we co-stained with segment-specific markers, Lotus Tetragonolobus Lectin (LTL)^97^,Tamm Horsfall protein (THP) antibody^98^, parvalbumin antibody^99^, and calbindin D28k antibody^99^, respectively. When assessing hyperplasia, we first selected tubular segments and counted DAPI-stained nuclei and PCNA fluorescence that overlapped with nuclei. We used seven images per sample and attempted to use a minimum of one PCNA count per sample. We quantified data from three microscope images per sample (20X setting) and combined values combined as the percentage of number of PCNA-stained nuclei to total number of nuclei counted (PCNA/nuclei × 100%).

To measure hypertrophy, we used immunohistochemistry (IHC) with quantitative morphometry to measure segment-specific cell size of vehicle- and furosemide-treated mice at 1 and 3 weeks. We performed quantitative morphometry for 30 tubular segments per sample using ImageJ software as previously described^100^. We defined cell area size by tubular segment area divided by total nuclei. We determined cell area by dividing the detected tubular area by the number of stained nuclei and compared these weighted values across cell types and time points as indicated.

### Flow Cytometry

We isolated kidney tubular cells from mice treated with vehicle or furosemide for 3 weeks. Once isolated and re-immersed in PBS buffer, we incubated samples at 1:200 dilutions with LTL-FITC to detect proximal tubular cells and Dolichos Biflorus Agglutinin–rhodamine (DBA-Rh) to detect principal cells^101,102^. During this incubation, we slowly rotated samples at room temperature before placing on ice. We added DAPI at a final concentration of 1 µM for live and dead cell discrimination. We then pooled samples in equal amounts for gating controls in all possible staining combinations. We analyzed stained cells on a Scanford (FACScan) flow cytometer at 100,000 events per sample. We systematically backgated to select for single cells, live cells, and lastly LTL-FITC or DBA-Rh fluorescently labeled cells. We performed data analysis using Flowjo (TreeStar Inc., Ashland, OR) and Prism GraphPad7 software (La Jolla, CA).

### Single Cell Isolation

We perfused anesthetized mice with cold PBS buffer and dissected whole kidney for cell isolation as previously described^102^. We minced samples 2X in modified HEPES Ringer buffer (mRING) using GentleMACs cell dissociator at h_cord setting at 1-minute durations. After spin down for 2 minutes at 1000 g, we resuspended pellets in mRING containing 0.2% collagenase and 0.2% hyaluronidase then shook at 2,600 rpm for 45 minutes at 37°C. We disrupted tubular fragments by passing the pellet gently through an 18-gauge needle 3x before 30-minute incubation with DNAse at 37°C. We further washed and centrifuged pellets at 1,000 rpm for 2 min multiple times until solution was clear in order to remove debris. For flow cytometry experiments, we re-suspended pellets in PBS, passed 2X through a 40 µm strainer and then 2X through a 20 µm strainer, spun one time at 1,000 g for 10 minutes, incubated with fluorophore-conjugated lectin and/or DAPI for 30 min at RT, and then placed on ice^102^. For single cell RNAseq experiments, we re-suspended pellets in 10 mL of patented modified PBS (mPBS) and passed through a 40 µm strainer right over a 20 µm filter and washed through with additional 10 mL mPBS. We then inverted the 20 µm filter over and washed a second time (10 mL mPBS) through a separate 20 µm filter to further increase yield [**Figure S3]**^102^.

For live/dead single cell analysis, we used the LIVE/DEAD™ Cell Vitality Assay Kit (ThermoFisher Scientific) per the manufacturer’s instructions. We used C_12_-resuzurin at 0.5 µM to detect live cells and 50 µM SYTOX Green to detect dead cells. To analyze the viability of isolated single cells, we gated 100,000 events for single cells, followed by Scanford Blue1 fluorescence detection for SYTOX Green (dead cells) and Yellow1 for Resorufin (end compound for live cells). Total live cells were any events within the population gated for Resorufin [**Figure S3b**].

### Single cell RNAseq using 500-gene targeted panel and BD™Rhapsody Platform

Since RNAseq analysis using a gene panel instead of whole transcriptome sequencing is advantageous for sequencing depth performance^103^, we developed a nephron-based 500-gene panel physiologic and pathophysiologic studies. This panel contained identity genes specific to cell types across the nephron, including segments 1 thru 3 of the proximal tubule (S1, S2, S3), thin descending limb, thin and thick ascending limb (TAL), distal convoluted tubule segment 1 (DCT1), DCT2, connecting tubule (CNT), principal cells of the cortical and medullary collecting duct (CCD-PCs, MCD-PCs), type A intercalated cells (α-ICs), and type B intercalated cells (β-ICs). We also included channels and ion transporters expressed throughout the nephron. We also included genes that demonstrated significant, segment-specific expression changes in our preliminary, whole transcriptome pilot study (data not shown). We also included genes associated with several cellular processes of interest, including but not limited to processes associated with tubular remodeling, e.g. hyperplasia, hypertrophy, IGF-1 signaling, ion transport, inflammation, and fibrosis [**Table S1**].

We dissociated an adult mouse kidney was dissociated into a single cell suspension and loaded into a BD Rhapsody cartridge for single cell mRNA capture on Rhapsody beads [**Figure S7**]. We retrieved beads from the cartridge and performed targeted panel amplification and sequencing library preparation followed by sequencing on a HiSeq Illumina platform. Across all six samples, we compared the number of targeted genes with a raw adjusted sequencing depth above four to define sequencing saturation (“pass genes”) and carried forward 451 pass genes out of our original 500 targeted panel [**Figure S8, Table 1**]. After removal of doublets(BD™algorithm^104^) and normalization of cell number across samples, we analyzed 22,154 total cells across six samples. We submitted this dataset to the Gene Expression Omnibus (http://www.ncbi.nlm.nih.gov/geo/).

### Computational Analysis of Single Cell RNASeq Data

We used the Python programming language (v3.6.4; http://python.org) along with the following core packages for all data processing, plotting, and analysis unless otherwise noted: SciPy (v1.0.0; http://scipy.org; https://doi.org/10.1109/MCSE.2011.36), Pandas (v0.23.4; http://pandas.pydata.org), Seaborn (v0.9.0; https://seaborn.pydata.org), and Matplotlib (v2.2.2; https://matplotlib.org)^105^. We removed genes whose counts were zero across all cells. For downstream clustering, dimensionality reduction, we added a pseudo-count of 1 to remaining genes and Log base 2-transformed the resulting counts.

To plot single-cell RNASeq data, we used Uniform Manifold Approximation and Projection^106^ to reduce the dimensionality of the log-transformed count data to two dimensions. We combined cells from vehicle- and furosemide-treated mice prior to the dimensionality reduction. Results shown are of combined dataset. To assess significant differences in numbers of cells per cluster between vehicle- and furosemide-treated mice we used negative binomial regression^107,108^ (StatsModels v0.8.0; http://statsmodels.org). For significant differences in cell proportions we used a generalized linear model with a Gaussian distribution. For both analyses, we adjusted for multiple comparisons using the Benjamini-Hochberg method^109^ and defined significance as a false discovery rate (FDR) < 0.1.

### Single-Cell RNASeq Differential Expression

To determine differentially expressed genes between treatment conditions, we used negative binomial regression^107,108^ per gene with terms for replicate, cell type, and treatment. As we were most interested in cell-type specific effects of furosemide, rather than global effects, we also included a cell type × treatment interaction term. We then corrected p-values for the coefficient corresponding to the interaction term using the Benjamini-Hochberg method^109^. To identify genes associated with cell types, we calculated the Cohen’s d effect size for each gene between cells inside a cluster and those outside of it and ranked genes by effect size.

### Western blotting

We homogenized whole kidney in buffer (250 mM sucrose and 10 mM triethanolamine, pH 7.6) containing protease inhibitors (1 mM phenylmethylsulfonyl fluoride (PMSF), 1 mM benzamidine, and 1× Complete Protease Inhibitor Mixture (Roche Applied Science, Penzberg, Germany) and phosphatase inhibitors (2 μM microcystin-LR, and 1× Phosphatase-Inhibitor Sets I and II (Calbiochem, Hayward, CA). We centrifuged homogenates at 4,000 g for 15 minutes to remove debris, then at 16,500 g for 60 minutes to obtain a membrane fraction. We determined protein concentration via the Lowry method (Bio-Rad DC protein assay, Hercules, CA) and generated immunoblots by SDS-PAGE. We quantified protein abundance by densitometry using ImageJ (NIH, Bethesda, MD).

### Generation of inducible tubule and collecting duct-specific IGF-1 receptor knockout mice

Professor Jens Bruning (University of Cologne) generously provided *Igf1r*^flox/flox^ mice, and we obtained Pax8-rtTA and TetOCre mice from the Jackson Laboratories (Bar Harbor, ME). Each strain was on a C57BL/6 background. Pax8-rtTA is expressed along the entire renal tubule and collecting system and is absent in vessels and glomeruli^110^. We bred *Igf1r*^flox/flox^ mice with either Pax8-rtTA or TetOCre mice, and then intercrossed the progeny with *Igf1r*^flox/flox^ mice to generate either *Igf1r*^Pax8^ or *Igf1r*^TetOCre^ mice [**Figure 9**]. We then interbred these lines to generate *Igf1r*^Pax8/TetOCre^ (iT-IGF1-RKO) mice. We used *Igf1r*^Pax8^ or *Igf1r*^TetOCre^ littermates of knockout mice as controls. We performed genotyping using previously published primers^110,111^ [**Table S4**]. To induce knockout of the IGF-1R gene, we fed mice older than 12 weeks 0.2 mg/mL doxycycline daily for 2 weeks as described previously^112^. We confirmed IGF-1 receptor deletion by immunoblotting of whole kidney lysate, as described above.

### Statistical Analysis

In experiments using four groups in a 2 by 2 scheme, we analyzed data by two-way ANOVA followed by the Sidak method to correct for multiple comparisons (Prism 6, San Diego, CA). For multiple pair-wise comparisons, we used a two-tailed t-test with Bonferroni correction. For histograms of weighted values, we used a Mann-Whitney non-parametric test. We defined significance as a two-tailed p-value < 0.05. The number of samples for each experiment is noted in the figure legends.

## Acknowledgements

The authors thank Glenn Chertow, Jonathan Maltzman, Arohan Subramanya, and Xiaoyi Zheng for helpful discussions. We thank Varsha Rao for assistance with library sequencing.

## REFERENCES

1. Buggey, J., et al. A reappraisal of loop diuretic choice in heart failure patients. Am Heart J 169, 323–333 (2015).

2. Hull, R.P. & Goldsmith, D.J. Nephrotic syndrome in adults. BMJ 336, 1185–1189 (2008).

3. Scaglione, S., et al. The Epidemiology of Cirrhosis in the United States: A Population-based Study. J Clin Gastroenterol 49, 690–696 (2015).

4. Khan, Y.H., Sarriff, A., Adnan, A.S., Khan, A.H. & Mallhi, T.H. Chronic Kidney Disease, Fluid Overload and Diuretics: A Complicated Triangle. PLoS One 11, e0159335 (2016).

5. Brater, D.C. Diuretic therapy. N Engl J Med 339, 387–395 (1998).

6. Seely, J.F. & Dirks, J.H. Site of action of diuretic drugs. Kidney Int 11, 1–8 (1977).

7. Sarafidis, P.A., Georgianos, P.I. & Lasaridis, A.N. Diuretics in clinical practice. Part I: mechanisms of action, pharmacological effects and clinical indications of diuretic compounds. Expert Opin Drug Saf 9, 243–257 (2010).

8. Cohen, N., et al. Serum magnesium aberrations in furosemide (frusemide) treated patients with congestive heart failure: pathophysiological correlates and prognostic evaluation. Heart 89, 411–416 (2003).

9. Ellison, D.H. Diuretic resistance: physiology and therapeutics. Semin Nephrol 19, 581–597 (1999).

10. Jentzer, J.C., DeWald, T.A. & Hernandez, A.F. Combination of loop diuretics with thiazide-type diuretics in heart failure. J Am Coll Cardiol 56, 1527–1534 (2010).

11. Kapelios, C.J., et al. High furosemide dose has detrimental effects on survival of patients with stable heart failure. Hellenic J Cardiol 56, 154–159 (2015).

12. Loon, N.R., Wilcox, C.S. & Unwin, R.J. Mechanism of impaired natriuretic response to furosemide during prolonged therapy. Kidney Int 36, 682–689 (1989).

13. Rao, V.S., et al. Compensatory Distal Reabsorption Drives Diuretic Resistance in Human Heart Failure. J Am Soc Nephrol 28, 3414–3424 (2017).

14. Ter Maaten, J.M., et al. Renal tubular resistance is the primary driver for loop diuretic resistance in acute heart failure. Eur J Heart Fail 19, 1014–1022 (2017).

15. Ellison, D.H., Velazquez, H. & Wright, F.S. Adaptation of the distal convoluted tubule of the rat. Structural and functional effects of dietary salt intake and chronic diuretic infusion. J Clin Invest 83, 113–126 (1989).

16. van Angelen, A.A., van der Kemp, A.W., Hoenderop, J.G. & Bindels, R.J. Increased expression of renal TRPM6 compensates for Mg(2+) wasting during furosemide treatment. Clin Kidney J 5, 535–544 (2012).

17. Kaissling, B., Bachmann, S. & Kriz, W. Structural adaptation of the distal convoluted tubule to prolonged furosemide treatment. Am J Physiol 248, F374–381 (1985).

18. Kaissling, B. & Stanton, B.A. Adaptation of distal tubule and collecting duct to increased sodium delivery. I. Ultrastructure. Am J Physiol 255, F1256–1268 (1988).

19. Loffing, J., et al. Thiazide treatment of rats provokes apoptosis in distal tubule cells. Kidney Int 50, 1180–1190 (1996).

20. Lalioti, M.D., et al. Wnk4 controls blood pressure and potassium homeostasis via regulation of mass and activity of the distal convoluted tubule. Nat Genet 38, 1124–1132 (2006).

21. Grimm, P.R., Coleman, R., Delpire, E. & Welling, P.A. Constitutively Active SPAK Causes Hyperkalemia by Activating NCC and Remodeling Distal Tubules. J Am Soc Nephrol 28, 2597–2606 (2017).

22. Lane, P.H. Furosemide treatment, angiotensin II, and renal growth and development in the rat. Pediatr Res 37, 747–754 (1995).

23. Lane, P.H., Tyler, L.D. & Schmitz, P.G. Chronic administration of furosemide augments renal weight and glomerular capillary pressure in normal rats. Am J Physiol 275, F230–234 (1998).

24. Kobayashi, S., Clemmons, D.R., Nogami, H., Roy, A.K. & Venkatachalam, M.A. Tubular hypertrophy due to work load induced by furosemide is associated with increases of IGF-1 and IGFBP-1. Kidney Int 47, 818–828 (1995).

25. Holzenberger, M., et al. IGF-1 receptor regulates lifespan and resistance to oxidative stress in mice. Nature 421, 182–187 (2003).

26. Chitnis, M.M., Yuen, J.S., Protheroe, A.S., Pollak, M. & Macaulay, V.M. The type 1 insulin-like growth factor receptor pathway. Clin Cancer Res 14, 6364–6370 (2008).

27. McCusker, R.H., Camacho-Hubner, C., Bayne, M.L., Cascieri, M.A. & Clemmons, D.R. Insulin-like growth factor (IGF) binding to human fibroblast and glioblastoma cells: the modulating effect of cell released IGF binding proteins (IGFBPs). J Cell Physiol 144, 244–253 (1990).

28. Segev, Y., et al. Renal hypertrophy in hyperglycemic non-obese diabetic mice is associated with persistent renal accumulation of insulin-like growth factor I. J Am Soc Nephrol 8, 436–444 (1997).

29. Lajara, R., et al. Dual regulation of insulin-like growth factor I expression during renal hypertrophy. Am J Physiol 257, F252–261 (1989).

30. Senthil, D., Choudhury, G.G., Abboud, H.E., Sonenberg, N. & Kasinath, B.S. Regulation of protein synthesis by IGF-I in proximal tubular epithelial cells. Am J Physiol Renal Physiol 283, F1226–1236 (2002).

31. Kiuchi-Saishin, Y., et al. Differential expression patterns of claudins, tight junction membrane proteins, in mouse nephron segments. J Am Soc Nephrol 13, 875–886 (2002).

32. Lee, J.W., Chou, C.L. & Knepper, M.A. Deep Sequencing in Microdissected Renal Tubules Identifies Nephron Segment-Specific Transcriptomes. J Am Soc Nephrol 26, 2669–2677 (2015).

33. Roy, A., Al-bataineh, M.M. & Pastor-Soler, N.M. Collecting duct intercalated cell function and regulation. Clin J Am Soc Nephrol 10, 305–324 (2015).

34. Trepiccione, F. & Capasso, G. SGK3: a novel regulator of renal phosphate transport? Kidney Int 80, 13–15 (2011).

35. Vallon, V., et al. SGLT2 mediates glucose reabsorption in the early proximal tubule. J Am Soc Nephrol 22, 104–112 (2011).

36. Woudenberg-Vrenken, T.E., Bindels, R.J. & Hoenderop, J.G. The role of transient receptor potential channels in kidney disease. Nat Rev Nephrol 5, 441–449 (2009).

37. Kriz, W. & Bankir, L. A standard nomenclature for structures of the kidney. The Renal Commission of the International Union of Physiological Sciences (IUPS). Kidney Int 33, 1–7 (1988).

38. Huang, Y., et al. TFRC promotes epithelial ovarian cancer cell proliferation and metastasis via up-regulation of AXIN2 expression. Am J Cancer Res 10, 131–147 (2020).

39. Chen, H.Y., et al. The protective role of Smad7 in diabetic kidney disease: mechanism and therapeutic potential. Diabetes 60, 590–601 (2011).

40. Kim, G.H. Renal effects of prostaglandins and cyclooxygenase-2 inhibitors. Electrolyte Blood Press 6, 35–41 (2008).

41. Palmer, B.F. Regulation of Potassium Homeostasis. Clin J Am Soc Nephrol 10, 1050–1060 (2015).

42. Roy, A., et al. Alternatively spliced proline-rich cassettes link WNK1 to aldosterone action. J Clin Invest 125, 3433–3448 (2015).

43. Ronzaud, C., et al. Renal tubular NEDD4-2 deficiency causes NCC-mediated salt-dependent hypertension. J Clin Invest 123, 657–665 (2013).

44. Al-Qusairi, L., et al. Renal Tubular Ubiquitin-Protein Ligase NEDD4-2 Is Required for Renal Adaptation during Long-Term Potassium Depletion. J Am Soc Nephrol 28, 2431–2442 (2017).

45. Shi, P.P., et al. Salt-sensitive hypertension and cardiac hypertrophy in mice deficient in the ubiquitin ligase Nedd4-2. Am J Physiol Renal Physiol 295, F462–470 (2008).

46. Debonneville, C., et al. Phosphorylation of Nedd4-2 by Sgk1 regulates epithelial Na(+) channel cell surface expression. EMBO J 20, 7052–7059 (2001).

47. Manning, J.A. & Kumar, S. Physiological Functions of Nedd4-2: Lessons from Knockout Mouse Models. Trends Biochem Sci 43, 635–647 (2018).

48. Andrew, M., et al. PAPPA2 as a Therapeutic Modulator of IGF-I Bioavailability: in Vivo and in Vitro Evidence. J Endocr Soc 2, 646–656 (2018).

49. Rogers, S.A., Powell-Braxton, L. & Hammerman, M.R. Insulin-like growth factor I regulates renal development in rodents. Dev Genet 24, 293–298 (1999).

50. Svensson, J., et al. Liver-derived IGF-I regulates kidney size, sodium reabsorption, and renal IGF-II expression. J Endocrinol 193, 359–366 (2007).

51. Hwa, V., Oh, Y. & Rosenfeld, R.G. The insulin-like growth factor-binding protein (IGFBP) superfamily. Endocr Rev 20, 761–787 (1999).

52. Bayascas, J.R. PDK1: the major transducer of PI 3-kinase actions. Curr Top Microbiol Immunol 346, 9–29 (2010).

53. Glass, D.J. PI3 kinase regulation of skeletal muscle hypertrophy and atrophy. Curr Top Microbiol Immunol 346, 267–278 (2010).

54. Rommel, C., et al. Mediation of IGF-1-induced skeletal myotube hypertrophy by PI(3)K/Akt/mTOR and PI(3)K/Akt/GSK3 pathways. Nat Cell Biol 3, 1009–1013 (2001).

55. Fayard, E., Xue, G., Parcellier, A., Bozulic, L. & Hemmings, B.A. Protein kinase B (PKB/Akt), a key mediator of the PI3K signaling pathway. Curr Top Microbiol Immunol 346, 31–56 (2010).

56. Mange, A., et al. FKBP4 connects mTORC2 and PI3K to activate the PDK1/Akt-dependent cell proliferation signaling in breast cancer. Theranostics 9, 7003–7015 (2019).

57. Spurlock, D.M., McDaneld, T.G. & McIntyre, L.M. Changes in skeletal muscle gene expression following clenbuterol administration. BMC Genomics 7, 320 (2006).

58. Brown, E.L., et al. PGC-1alpha and PGC-1beta Increase Protein Synthesis via ERRalpha in C2C12 Myotubes. Front Physiol 9, 1336 (2018).

59. Hemmings, B.A. & Restuccia, D.F. PI3K-PKB/Akt pathway. Cold Spring Harb Perspect Biol 4, a011189 (2012).

60. Packer, L., et al. Osteopontin is a downstream effector of the PI3-kinase pathway in melanomas that is inversely correlated with functional PTEN. Carcinogenesis 27, 1778–1786 (2006).

61. Junaid, A. & Amara, F.M. Osteopontin: correlation with interstitial fibrosis in human diabetic kidney and PI3-kinase-mediated enhancement of expression by glucose in human proximal tubular epithelial cells. Histopathology 44, 136–146 (2004).

62. Lin, Y.H. & Yang-Yen, H.F. The osteopontin-CD44 survival signal involves activation of the phosphatidylinositol 3-kinase/Akt signaling pathway. J Biol Chem 276, 46024–46030 (2001).

63. The Ambulatory Care Drug Database System. (2015).

64. Alexander, R.T. & Dimke, H. Effect of diuretics on renal tubular transport of calcium and magnesium. Am J Physiol Renal Physiol 312, F998–F1015 (2017).

65. Rabkin, R. & Fervenza, F.C. Renal hypertrophy and kidney disease in diabetes. Diabetes Metab Rev 12, 217–241 (1996).

66. Fervenza, F.C., Tsao, T., Hsu, F. & Rabkin, R. Intrarenal insulin-like growth factor-1 axis after unilateral nephrectomy in rat. J Am Soc Nephrol 10, 43–50 (1999).

67. Tsao, T., et al. Expression of insulin-like growth factor-I and transforming growth factor-beta in hypokalemic nephropathy in the rat. Kidney Int 59, 96–105 (2001).

68. Wolf, G., et al. Angiotensin II-induced hypertrophy of proximal tubular cells requires p27Kip1. Kidney Int 64, 71–81 (2003).

69. Wolf, G. & Neilson, E.G. Angiotensin II induces cellular hypertrophy in cultured murine proximal tubular cells. Am J Physiol 259, F768–777 (1990).

70. Liu, B. & Preisig, P.A. Compensatory renal hypertrophy is mediated by a cell cycle-dependent mechanism. Kidney Int 62, 1650–1658 (2002).

71. Christensen, S. & Petersen, J.S. Effects of furosemide on renal haemodynamics and proximal tubular sodium reabsorption in conscious rats. Br J Pharmacol 95, 353–360 (1988).

72. Modena, B., et al. Furosemide stimulates renin expression in the kidneys of salt-supplemented rats. Pflugers Arch 424, 403–409 (1993).

73. Stanton, B.A. & Kaissling, B. Regulation of renal ion transport and cell growth by sodium. Am J Physiol 257, F1–10 (1989).

74. Brater, D.C. Diuretic resistance: mechanisms and therapeutic strategies. Cardiology 84 Suppl 2, 57–67 (1994).

75. Wheatley, S.P. & Altieri, D.C. Survivin at a glance. J Cell Sci 132(2019).

76. Delaney, J.A., Levesque, L.E., Etminan, M. & Suissa, S. Furosemide use and hospitalization for benign prostatic hyperplasia. Can J Clin Pharmacol 13, e75–80 (2006).

77. Shiozaki, A., et al. Furosemide, a blocker of Na+/K+/2Cl- cotransporter, diminishes proliferation of poorly differentiated human gastric cancer cells by affecting G0/G1 state. J Physiol Sci 56, 401–406 (2006).

78. Han, D.C., Hoffman, B.B., Hong, S.W., Guo, J. & Ziyadeh, F.N. Therapy with antisense TGF-beta1 oligodeoxynucleotides reduces kidney weight and matrix mRNAs in diabetic mice. Am J Physiol Renal Physiol 278, F628–634 (2000).

79. Vallon, V. The proximal tubule in the pathophysiology of the diabetic kidney. Am J Physiol Regul Integr Comp Physiol 300, R1009–1022 (2011).

80. Huang, H.C. & Preisig, P.A. G1 kinases and transforming growth factor-beta signaling are associated with a growth pattern switch in diabetes-induced renal growth. Kidney Int 58, 162–172 (2000).

81. Abdallah, J.G., et al. Loop diuretic infusion increases thiazide-sensitive Na(+)/Cl(-)- cotransporter abundance: role of aldosterone. J Am Soc Nephrol 12, 1335–1341 (2001).

82. Flyvbjerg, A., et al. The role of growth hormone, insulin-like growth factors (IGFs), and IGF-binding proteins in experimental diabetic kidney disease. Metabolism 44, 67–71 (1995).

83. Flyvbjerg, A., et al. Insulin-like growth factor I in initial renal hypertrophy in potassium-depleted rats. Am J Physiol 262, F1023–1031 (1992).

84. Chen, J.K., et al. Phosphatidylinositol 3-kinase signaling determines kidney size. J Clin Invest 125, 2429–2444 (2015).

85. Bach, L.A. & Hale, L.J. Insulin-like growth factors and kidney disease. Am J Kidney Dis 65, 327–336 (2015).

86. Miller, S.B., Martin, D.R., Kissane, J. & Hammerman, M.R. Insulin-like growth factor I accelerates recovery from ischemic acute tubular necrosis in the rat. Proc Natl Acad Sci U S A 89, 11876–11880 (1992).

87. Li, L., et al. Absence of renal enlargement in fructose-fed proximal-tubule-select insulin receptor (IR), insulin-like-growth factor receptor (IGF1R) double knockout mice. Physiol Rep 4(2016).

88. Jones, J.I., D’Ercole, A.J., Camacho-Hubner, C. & Clemmons, D.R. Phosphorylation of insulin-like growth factor (IGF)-binding protein 1 in cell culture and in vivo: effects on affinity for IGF-I. Proc Natl Acad Sci U S A 88, 7481–7485 (1991).

89. Salih, D.A., et al. Insulin-like growth factor-binding protein 5 (Igfbp5) compromises survival, growth, muscle development, and fertility in mice. Proc Natl Acad Sci U S A 101, 4314–4319 (2004).

90. Kalus, W., et al. Structure of the IGF-binding domain of the insulin-like growth factor-binding protein-5 (IGFBP-5): implications for IGF and IGF-I receptor interactions. EMBO J 17, 6558–6572 (1998).

91. Tripathi, G., et al. IGF-independent effects of insulin-like growth factor binding protein-5 (Igfbp5) in vivo. FASEB J 23, 2616–2626 (2009).

92. Ding, M., Bruick, R.K. & Yu, Y. Secreted IGFBP5 mediates mTORC1-dependent feedback inhibition of IGF-1 signalling. Nat Cell Biol 18, 319–327 (2016).

93. O’Neill, B.T., et al. Insulin and IGF-1 receptors regulate FoxO-mediated signaling in muscle proteostasis. J Clin Invest 126, 3433–3446 (2016).

94. Nizar, J.M., Bouby, N., Bankir, L. & Bhalla, V. Improved protocols for the study of urinary electrolyte excretion and blood pressure in rodents: use of gel food and stepwise changes in diet composition. Am J Physiol Renal Physiol (2018).

95. Hall, P.A., et al. Proliferating cell nuclear antigen (PCNA) immunolocalization in paraffin sections: an index of cell proliferation with evidence of deregulated expression in some neoplasms. J Pathol 162, 285–294 (1990).

96. Xie, C., et al. Down-regulated CFTR During Aging Contributes to Benign Prostatic Hyperplasia. J Cell Physiol 230, 1906–1915 (2015).

97. Laitinen, L., Virtanen, I. & Saxen, L. Changes in the glycosylation pattern during embryonic development of mouse kidney as revealed with lectin conjugates. J Histochem Cytochem 35, 55–65 (1987).

98. Stricklett, P.K., Taylor, D., Nelson, R.D. & Kohan, D.E. Thick ascending limb-specific expression of Cre recombinase. Am J Physiol Renal Physiol 285, F33–39 (2003).

99. Loffing, J., et al. Distribution of transcellular calcium and sodium transport pathways along mouse distal nephron. Am J Physiol Renal Physiol 281, F1021–1027 (2001).

100. Chang, M., et al. Changes in cell-cycle kinetics responsible for limiting somatic growth in mice. Pediatr Res 64, 240–245 (2008).

101. Holthofer, H., Schulte, B.A. & Spicer, S.S. Expression of binding sites for Dolichos biflorus agglutinin at the apical aspect of collecting duct cells in rat kidney. Cell Tissue Res 249, 481–485 (1987).

102. Labarca, M., et al. Harvest and primary culture of the murine aldosterone-sensitive distal nephron. Am J Physiol Renal Physiol 308, F1306–1315 (2015).

103. Wooderchak-Donahue, W.L., et al. A direct comparison of next generation sequencing enrichment methods using an aortopathy gene panel- clinical diagnostics perspective. BMC Med Genomics 5, 50 (2012).

104. BD Single Cell Genomics Bioinformatics Handbook. Vol. 54169 (ed. Biosciences, B.) 98–99 (Becton, Dickinson, and Company, San Jose, CA 95131, 2018).

105. Hunter, J.D. Matplotlib: A 2D graphics environment. Comput Sci Eng 9, 90–95 (2007).

106. Becht, E., et al. Dimensionality reduction for visualizing single-cell data using UMAP. Nat Biotechnol (2018).

107. Love, M.I., Huber, W. & Anders, S. Moderated estimation of fold change and dispersion for RNA-seq data with DESeq2. Genome Biol 15, 550 (2014).

108. Stuart, T., et al. Comprehensive Integration of Single-Cell Data. Cell 177, 1888–1902 e1821 (2019).

109. Benjamini, Y. & Hochberg, Y. Controlling the False Discovery Rate - a Practical and Powerful Approach to Multiple Testing. J R Stat Soc B 57, 289–300 (1995).

110. Traykova-Brauch, M., et al. An efficient and versatile system for acute and chronic modulation of renal tubular function in transgenic mice. Nat Med 14, 979–984 (2008).

111. Kloting, N., et al. Autocrine IGF-1 action in adipocytes controls systemic IGF-1 concentrations and growth. Diabetes 57, 2074–2082 (2008).

112. Nizar, J.M., Shepard, B.D., Vo, V.T. & Bhalla, V. Renal tubule insulin receptor modestly promotes elevated blood pressure and markedly stimulates glucose reabsorption. JCI Insight 3(2018).

