## Supplementary Figure Legends and Tables for "Proximal and Distal Nephron-specific Adaptation to Furosemide"

**Supplementary Figure S1: Treatment of mice with vehicle vs. furosemide.** (a) Experimental protocol for one and three week diuretic treatments. (b) There was no detectable difference in mouse weight between mice fed vehicle vs. furosemide diet for one week (*left panel*, N=5 mice /group) or three weeks (*right panel*, N=26-27 mice/group). (c) One-week analysis shows gel food intake was similar between groups whereas water intake was significantly higher in mice fed furosemide diet (N=7-8 mice/group). For one- and three-week treatment experiments, the efficacy of diuretic was monitored by randomly choosing 2-3 mice per group to compare water intake. (d) Select window within 3-week treatment regimen demonstrating higher water intake for furosemide-fed mice (N=3 mice/group). Data are presented as mean ± SEM. *p-value < 0.05 vs. vehicle treatment.

**Supplementary Figure S2: Furosemide-induced proximal and distal tubular hypertrophy is observed at the single cell level using flow cytometry analysis.** (a) Gating of forward and forward width scatter for single cells, DAPI-free for viable cells, and either proximal tubule cells bound to LTL-FITC (y-axis) or CNT/CCD principal cells bound to DBA-rhodamine (x-axis). (b) Flow cytometry analysis of PT cells (*left panel*) and CNT/CCD principal cells (*right panel*) normalized to 3-week vehicle mean-averaged values (N=5-6 mice/group). Data are presented as mean ± SEM. *p-value < 0.05 vs. vehicle treatment.

**Supplementary Figure S3: Isolation of Single Viable Kidney Cells for single cell RNAseq.** (a) Protocol for single cell digestion for RNAseq. Kidneys were perfused and digested followed by mincing and trituration before loading on sequencing cartridges. (b) To test for digestion quality and viability, in parallel, digested single cells were gated (*left panel*, FSC to FSCW), followed by gating for fluorescent metabolic end product $C_{12}$-resorufin for live cells (*right panel*).

**Supplementary Figure S4: Absence of batch effect across samples.** UMAP projection for each individual sample from three vehicle- (a) and furosemide-treated (b) mice.

**Supplementary Figure S5: Larger and smaller proportions of distinct cell populations with one week furosemide treatment.** Proportion of total cells per cluster (segment/cell type) in vehicle- (*purple*) and furosemide-treated (*green*) mice. Each dot represents one mouse. Data is represented by violin plots. *, indicates cell clusters with false discovery rate < 0.1. Glom, glomerular cells; S1/S2; Proximal tubule subsegments S1/S2, S2/S3; Proximal tubule subsegments S2/S3; TAL, thick ascending limb; DCT1, distal convoluted tubule segment 1; DCT2, distal convoluted tubule segment 2; CNT, connecting tubule; CCD PCs, principal cells of the cortical collecting duct; Alpha ICs, type A intercalated cells; Beta ICs, type B intercalated cells; MCD PCs, principal cells of the outer medullary collecting duct; Immune cells, lymphocytes and macrophages; Cluster X, *Wnt4*(+) cells. N=3 mice/group.

**Supplementary Figure S6: Expression of IGF-1 signaling genes in vehicle and furosemide-treated mice.** Representative immunoblot (*top panel*) and densitometry (*bottom panel*) of whole kidney lysates probed for IGF-1 after three weeks in vehicle- (*purple*) vs. furosemide-fed (*green*) mice. *, non-specific band. IGF-1 relative to non-specific band in vehicle- (*purple*) vs. furosemide-fed (*green*) mice after three weeks. N=4 mice/group. Data are normalized to vehicle treatment and presented as mean ± SEM. *p-value < 0.05 vs. vehicle treatment.

**Supplementary Figure S7: Utilization of BD Rhapsody for single cell capture and library preparation of a dissociated mouse kidney.** (a) We loaded isolated single kidney cells from vehicle- and furosemide-treated mice into a Rhapsody cartridge. Rhapsody mRNA capture beads were loaded to saturation and cells were lysed. Following mRNA hybridization, Rhapsody beads were retrieved and subjected to cDNA synthesis and sequencing library preparation. (b) Schematic of Rhapsody capture beads loaded with mRNA and final sequencing library products from Rhapsody library preparation. Univ = Universal sequence, MI = Molecular index, CL = Cell label, dT = oligo(dT).

**Supplementary Figure S8: Quality control across single cell RNAseq samples.** Undetectable (*black*), low depth (*gray*), pass genes (*white*) from the 500-gene targeted panel across each of the six samples (N=3 mice/group). Vehicle-treated mice are 1,3,5 (*purple*) and furosemide-treated mice are 2,4,6 (*green*).

**Supplementary Table S1: Differentially regulated genes within cell clusters from vehicle- vs. furosemide-treated mice**

See table below

z-score | FDR

| Gene | Glomeruli | | S1/S2 | | S1/S2/S3 | | S2/S3 | | Thin Descending Limb | | TAL | | DCT1 | | DCT2 | | CNT | | CCD PCs | | Alpha ICs | | Beta ICs | | MCD PCs | | Immune Cells | | Cluster X | |
|---|---|---|---|---|---|---|---|---|---|---|---|---|---|---|---|---|---|---|---|---|---|---|---|---|---|---|---|---|---|---|
| Acy3 | | | | | | | | | | | -0.338134 | 2.79E-14 | | | | | | | | | | | | | | | | | | |
| Adamts1 | | | | | -0.274325 | 0.0064714 | | | | | | | | | | | -0.717498 | 6.91E-08 | -0.512849 | 5.19E-10 | | | | | | | | | | |
| Adh1 | | | 0.191519 | 0.0000332 | | | -0.16248 | 0.0000061 | | | | | | | | | | | | | | | | | | | | | | |
| Adora2b | | | | | | | | | | | -0.964312 | 0.00098 | | | | | | | | | | | | | | | | | | |
| Ahsa1 | | | | | | | | | | | | | | | | | 0.3415658 | 0.0005751 | | | | | | | | | | | | |
| Akr1b3 | | | | | | | | | -1.022728 | 0.0050011 | | | | | | | | | | | | | | | | | | | | |
| Akt1 | | | | | | | | | | | | | | | | | 0.2623537 | 0.0069684 | | | | | | | | | 0.4868756 | 0.0031817 | | |
| Aldob | | | | | -0.066894 | 0.0082322 | | | | | | | | | | | | | | | | | | | | | | | | |
| Apoe | | | | | | | | | | | -1.240407 | 0.0003195 | | | | | | | | | | | | | | | | | | |
| Aqp1 | | | | | -0.13907 | 0.0017146 | -0.107064 | 0.0021113 | -0.639823 | 2.11E-09 | | | | | | | | | | | | | | | | | | | | |
| Atf3 | | | | | | | 0.2510248 | 0.0004895 | -0.232417 | 0.0028175 | -0.30117 | 1.22E-11 | | | | | | | | | | | | | | | | | | |
| Atf4 | | | | | | | | | 0.2659425 | 0.0006207 | -0.204774 | 1.19E-08 | | | | | | | | | | | | | | | | | | |
| Atf6 | | | | | | | -0.258791 | 0.007471 | | | | | | | | | | | | | | | | | | | | | | |
| Atp1a1 | | | 0.0966283 | 0.0026872 | | | | | -0.296669 | 9.89E-06 | -0.227091 | 3.14E-30 | 0.1340866 | 0.0000151 | | | 0.4769239 | 1.07E-25 | 0.3255963 | 5.05E-14 | | | 0.1708457 | 0.0026633 | 0.3587348 | 0.0003018 | | | | |
| Atp1b1 | | | | | -0.160661 | 4.45E-08 | -0.20212 | 1.01E-10 | -0.611616 | 6.2E-17 | | | 0.1329097 | 0.0003688 | | | 0.2111119 | 0.0000258 | | | | | | | | | | | | |
| Atp2a2 | | | -0.172741 | 0.0067103 | -0.356913 | 1.42E-17 | -0.235741 | 0.000169 | | | -0.141449 | 0.0024101 | | | | | | | | | | | | | | | | | | |
| Atp4a | | | | | | | | | | | 0.4674786 | 1.57E-17 | -0.38916 | 4.73E-13 | | | | | 0.6679984 | 8.55E-13 | | | 0.6865087 | 2.59E-09 | 0.7856246 | 0.0005783 | | | | |
| Atp6ap2 | | | -0.110698 | 0.0006734 | -0.113841 | 0.0002781 | -0.166982 | 1.13E-08 | -0.303525 | 0.0005252 | -0.122207 | 0.0000548 | | | | | 0.2124549 | 0.0003648 | | | | | | | | | | | | |
| Atp6v1c2 | | | | | | | | | -1.544718 | 0.0085505 | | | 0.6254742 | 1.31E-06 | | | | | -0.980662 | 0.0025212 | | | | | | | | | | |
| Atp6v1g3 | | | | | | | | | | | | | | | | | | | | | | | 0.2808744 | 5.7E-08 | 0.319453 | 0.0002202 | | | | |
| Bak1 | | | | | | | | | | | | | 0.2417121 | 0.0041406 | | | 0.5527643 | 9.97E-08 | | | | | | | | | | | | |
| Bax | | | | | | | 0.1870558 | 0.0000393 | | | | | | | | | | | | | | | | | | | 0.6156459 | 0.000098 | | |
| Bcat1 | | | | | | | | | -0.559455 | 0.0000148 | | | | | | | | | -1.333151 | 0.0012643 | | | | | | | | | -1.173733 | 0.0000013 |
| Becn1 | | | | | | | | | | | | | | | | | 0.4214349 | 3.06E-10 | | | | | | | | | | | | |
| Bhmt | | | | | | | 0.5475964 | 0.0062956 | | | | | | | | | | | | | | | | | | | | | | |
| Birc5 | | | | | | | | | | | | | 2.7026967 | 0.0052187 | | | | | | | | | | | | | | | | |
| Bok | | | | | | | -0.224255 | 0.0021371 | -0.958149 | 1.36E-08 | | | | | | | | | | | | | | | | | | | | |
| C1galt1 | | | | | | | | | | | -0.24412 | 5.28E-09 | | | | | | | | | | | | | | | | | | |
| C1galt1c1 | | | | | | | | | | | | | | | | | 0.3025008 | 0.0009796 | | | | | | | 0.387513 | 0.0020698 | | | | |
| C3 | | | | | | | | | | | | | | | | | | | | | | | | | | | 1.3350601 | 0.0059051 | | |
| Calb1 | | | 1.0861556 | 9.65E-07 | 1.7291654 | 0.0019289 | 0.9850262 | 8.17E-08 | | | 1.1217886 | 4.47E-07 | 0.4482796 | 3.04E-17 | | | 1.3124922 | 7.54E-82 | 0.7137094 | 5.43E-17 | 1.2271233 | 0.0025522 | 0.9402249 | 2.95E-07 | | | | | | |
| Calm1 | | | | | | | | | | | 0.0966782 | 0.0001055 | | | | | 0.2232028 | 0.0000993 | -0.180553 | 0.0010036 | | | | | | | 0.3075031 | 0.0041412 | | |
| Casp3 | | | | | | | | | | | -0.222729 | 0.0001147 | | | | | 0.4105612 | 0.0000326 | | | | | 0.3370822 | 0.0015146 | 0.3960038 | 0.0091674 | | | | |
| Casr | | | | | | | | | | | -0.283161 | 6.17E-09 | | | | | | | | | | | | | | | | | | |
| Ccl2 | | | | | | | | | | | | | | | | | | | | | | | | | | | 0.8614864 | 0.0000136 | | |
| Ccnd1 | | | -0.421554 | 6.57E-12 | -0.554833 | 1.81E-17 | -0.388296 | 0.000831 | -0.478312 | 0.0091154 | -0.991842 | 1.08E-151 | | | | | | | -0.20967 | 0.0036869 | | | | | | | | | | |
| Cd8a | | | | | | | | | | | | | | | | | 0.9609729 | 0.0000556 | | | | | | | | | | | | |
| Cdh1 | | | | | | | | | | | -0.087255 | 0.0022613 | | | | | | | | | | | | | | | | | | |
| Cdk9 | | | | | | | | | | | 0.1812173 | 0.0000171 | | | | | | | | | | | | | | | | | | |
| Cdkn1a | | | | | | | 0.5421712 | 0.0082203 | | | | | | | | | | | | | | | | | | | | | | |
| Cdkn1b | | | | | -0.310661 | 0.000102 | -0.355768 | 0.0009528 | -0.609236 | 0.0009429 | -0.201433 | 0.0001277 | | | | | | | | | | | | | | | | | | |
| Cela1 | | | | | | | 0.7842251 | 0.0086916 | | | | | | | | | | | | | | | | | | | | | | |
| Ckb | | | | | | | | | | | -0.442821 | 3.98E-56 | 0.3206938 | 1.5E-13 | 1.3102781 | 2.55E-15 | 1.334707 | 1.45E-07 | | | | | | | 1.0109736 | 6.44E-14 | 0.5117939 | 0.0017164 | | |
| Clcnka | | | | | | | -0.701797 | 0.0021371 | -1.320533 | 4.39E-06 | -0.468234 | 8.65E-64 | 0.1162451 | 0.0075398 | | | 0.5896289 | 0.000831 | | | | | 0.4274679 | 0.0000863 | | | | | | |
| Clcnkb | | | | | | | | | | | 0.5202181 | 2.42E-64 | -0.100439 | 0.004889 | | | 0.1484045 | 0.0005266 | 0.178434 | 0.0055069 | | | | | 0.3753774 | 3.66E-08 | | | | |
| Cldn10 | | | | | | | | | | | -0.188285 | 2.27E-09 | | | | | | | | | | | | | | | | | | |
| Cldn16 | | | | | | | | | | | 0.6363913 | 2.09E-58 | | | | | | | | | | | | | | | | | | |
| Cldn4 | | | | | | | | | | | | | | | | | | | | | | | | | | | | | 1.1532292 | 0.0003552 |
| Clu | | | 0.7520925 | 0.0002672 | | | | | 0.860355 | 0.00002 | 1.0106772 | 6.11E-07 | | | | | 0.6560188 | 6.94E-24 | | | | | | | | | | | 1.4130588 | 0.0068269 |
| Col1a1 | 1.7439158 | 0.0076568 | | | | | | | | | | | | | | | | | | | | | | | | | | | | |
| Col4a1 | | | | | | | | | 0.6414166 | 3.24E-11 | | | 0.5078287 | 0.0048935 | | | | | | | | | | | | | | | | |
| Col4a3 | | | | | -0.300213 | 5.31E-08 | -0.284345 | 0.0000305 | -0.430829 | 0.0032415 | | | | | | | 0.336389 | 0.0020352 | -0.351372 | 0.0062944 | | | | | -0.707895 | 0.0072828 | | | | |
| Col4a4 | | | | | -0.223939 | 0.0013698 | -0.251597 | 0.002124 | | | | | -0.195713 | 0.0034167 | | | | | | | | | | | | | | | | |
| Cops5 | | | | | | | | | | | | | | | | | 0.3946113 | 1.04E-09 | | | | | 0.2415366 | 0.0005116 | 0.3201779 | 0.0010795 | | | | |
| Creb3 | | | | | | | | | | | 0.1483104 | 0.0083486 | | | | | | | | | | | | | | | | | | |
| Crem | | | | | | | | | | | | | | | | | | | -0.468732 | 3.78E-09 | | | | | | | | | | |
| Cryab | | | | | | | | | | | -0.476537 | 0.0017157 | | | | | 0.3433096 | 0.0074138 | | | | | | | | | | | | |
| Ctxn3 | | | | | | | -0.679863 | 9.68E-07 | | | | | | | | | | | | | | | | | | | | | | |
| Cul3 | | | | | | | -0.206598 | 0.0006076 | | | | | | | | | | | | | | | | | | | | | | |
| Cux1 | | | | | | | | | | | 0.2243699 | 1.84E-12 | | | | | | | 0.2329151 | 0.0019434 | | | | | | | | | | |
| Cxcl1 | | | | | -0.384403 | 2.54E-06 | 0.3225153 | 1.31E-06 | | | | | | | | | | | | | | | | | | | | | | |
| Cxcl10 | | | | | | | 0.29303 | 0.0090856 | | | 0.4787725 | 2.56E-07 | | | | | | | | | | | | | | | | | | |
| Cyp1a1 | | | | | | | | | | | | | | | | | | | | | | | 0.479879 | 0.0033034 | | | | | | |
| Cyp2a4 | | | 0.3548319 | 0.0000218 | | | 0.2671907 | 0.00034 | | | | | | | | | | | | | | | | | | | | | | |
| Cyp4b1 | 0.6473321 | 0.0004243 | | | | | 0.1700693 | 0.0000277 | | | | | | | | | | | | | | | | | | | | | | |
| Cyr61 | -0.716333 | 0.0086794 | 0.2381341 | 0.0009291 | | | | | | | -0.645705 | 4.13E-22 | | | | | | | | | | | | | | | | | | |
| Dnajb1 | | | -0.263618 | 0.0024866 | -0.420538 | 8.55E-13 | -0.494399 | 1.52E-13 | | | -0.37415 | 1.93E-09 | | | | | | | | | | | | | | | | | | |
| Edem3 | | | | | -0.235181 | 0.0003406 | -0.271568 | 0.0004673 | | | | | | | | | | | | | | | | | | | | | | |
| Edn1 | -0.827058 | 0.0000168 | | | | | | | | | | | | | | | | | | | | | | | | | | | | |
| Egf | | | | | | | | | | | 0.1781324 | 1.59E-11 | -0.232965 | 3.41E-09 | | | | | | | | | | | | | | | | |
| Egln2 | | | | | | | | | | | | | | | | | 0.4375176 | 0.0000714 | | | | | | | | | 0.7217904 | 0.0026833 | | |
| Egr1 | | | | | -0.311145 | 2.65E-07 | | | | | -0.462631 | 7.06E-28 | | | | | | | | | | | | | | | | | | |
| Eif2ak3 | | | | | | | | | | | -0.189158 | 0.0005726 | | | | | | | | | | | | | | | | | | |
| Eif2b4 | | | | | | | | | | | | | | | | | 0.3869176 | 0.0030973 | | | | | | | 0.5713567 | 0.0053459 | | | | |
| Elk1 | | | | | | | | | | | 0.4646489 | 0.0010155 | | | | | | | | | | | | | | | | | | |
| Emp1 | | | | | | | | | | | | | -0.428518 | 0.0020352 | | | | | -0.597824 | 5.78E-23 | | | | | | | | | | |
| Fabp1 | | | 1.0299678 | 0.0004111 | | | | | | | | | | | | | | | | | | | | | | | | | | |
| Fas | | | | | | | | | | | -0.478727 | 0.0002067 | | | | | | | | | | | | | | | | | | |
| Fkbp1a | | | | | | | | | -0.239185 | 0.0027135 | | | | | | | 0.2387649 | 0.0002568 | | | | | | | | | | | | |
| Fkbp4 | | | | | | | | | | | | | | | | | 0.3574168 | 3.18E-06 | 0.2042423 | 0.0056281 | | | | | | | | | | |
| Fos | | | | | -0.186612 | 1.42E-06 | | | -0.258327 | 0.0037269 | -0.147243 | 0.0000656 | -0.123833 | 0.0046101 | | | | | 0.2473613 | 0.0053077 | | | | | | | | | | |
| Foxo3 | | | | | | | | | | | -0.295146 | 7.79E-13 | -0.394972 | 1.59E-10 | | | | | | | | | | | | | | | | |
| Fst | | | | | | | | | -1.803319 | 2.56E-08 | | | | | | | | | | | | | | | | | | | | |
| Fxyd2 | | | | | | | | | -2.688678 | 6.88E-07 | -0.93825 | 1.3E-55 | 0.1730003 | 0.0006292 | 0.4739339 | 0.0055202 | | | | | | | 0.251146 | 0.0069684 | | | | | | |
| Fxyd4 | | | | | | | | | | | | | | | | | -0.395313 | 1.83E-07 | -0.27224 | 4.22E-07 | | | | | | | | | | |
| G6pc | | | 0.3668018 | 3.36E-18 | 0.2893118 | 8.22E-11 | 0.2748081 | 2.37E-07 | | | -1.319744 | 2.92E-41 | | | | | | | | | | | | | | | | | | |
| Gpd1 | | | | | | | | | | | | | | | | | | | | | | | 0.3282171 | 0.0000681 | | | | | | |
| Gprc5a | | | | | | | | | | | | | | | | | | | -0.771117 | 0.0026872 | | | | | | | | | | |
| Gpx1 | | | 0.1288503 | 0.0000207 | 0.1368336 | 6.54E-07 | 0.2181613 | 3.03E-18 | -0.336345 | 0.0000266 | | | 0.1682729 | 1.95E-07 | | | 0.4515559 | 1.78E-20 | | | 0.3310493 | 0.0032869 | 0.494328 | 4.31E-17 | | | 0.3590385 | 0.0003746 | | |
| Gpx6 | | | | | | | | | | | -0.716896 | 7.59E-141 | 0.3801428 | 2.36E-10 | 1.0493439 | 1.54E-11 | | | | | | | | | | | | | | |
| Guca2a | | | | | | | | | | | | | | | | | | | | | | | | | 0.533269 | 0.0039233 | | | | |
| Hbegf | | | | | | | | | | | -0.419173 | 2.13E-07 | -0.52878 | 8.22E-08 | | | | | -0.448716 | 2.87E-11 | | | -0.444086 | 0.0007743 | | | | | | |
| Hdac2 | | | | | | | | | | | | | | | | | | | | | | | | | 0.370098 | 0.0053415 | | | | |
| Hif1a | | | | | | | | | -0.435718 | 0.0001708 | -0.134654 | 0.0045525 | | | | | | | | | | | 0.2837722 | 0.0057739 | | | | | | |
| Hist1h1c | -0.532274 | 0.0014762 | | | | | | | -1.134587 | 1.22E-27 | -1.330787 | 7.65E-70 | | | | | | | -0.599072 | 3.25E-07 | | | | | -0.576261 | 0.0034443 | | | | |
| Hmox1 | | | 0.6065448 | 9.55E-17 | 0.5024364 | 2.48E-15 | 0.5685873 | 3.39E-17 | 1.4211998 | 1.04E-09 | | | | | 1.1728297 | 0.0045525 | | | | | | | 0.5153059 | 0.0034167 | | | | | | |
| Hnf1a | | | | | | | | | | | -0.400842 | 0.0015593 | | | | | | | | | | | | | | | | | | |
| Hnf1b | | | -0.202798 | 0.0023908 | -0.302623 | 1.96E-10 | -0.311073 | 4.14E-09 | | | | | | | | | | | | | | | | | | | | | | |
| Hsd11b2 | | | | | 0.7265978 | 0.0003018 | | | | | 1.0478104 | 0.0003378 | | | | | 0.2072319 | 0.0000111 | 0.192887 | 5.16E-08 | | | | | | | | | 1.4790822 | 0.0000701 |
| Hspa1a | -0.528805 | 0.0003166 | | | -1.070839 | 2.79E-14 | -1.344334 | 8.43E-08 | | | -0.235229 | 0.0014266 | -0.896615 | 2.51E-18 | | | | | | | | | -0.761827 | 2.12E-07 | | | | | | |
| Hspa5 | | | -0.539784 | 4.58E-33 | -0.602557 | 3.97E-51 | -0.356506 | 4.95E-15 | | | 0.1100844 | 0.001822 | 0.3066861 | 4.2E-10 | | | 0.4982194 | 1.89E-14 | | | | | -0.418983 | 2.13E-07 | | | | | | |
| Icam1 | | | 0.4552821 | 9.35E-08 | | | 0.3030613 | 0.0000584 | 0.6738592 | 0.0019943 | | | | | | | | | | | | | | | | | | | | |
| Id3 | -0.42123 | 0.0009224 | -0.502452 | 2.63E-22 | -0.496638 | 6.46E-25 | -0.579435 | 1.63E-10 | | | | | -0.481225 | 1.56E-11 | -0.520462 | 0.0008364 | -0.247392 | 0.009251 | | | | | | | | | | | | |
| Ifit1 | | | | | | | | | | | 1.2537162 | 0.0015188 | | | | | | | | | | | | | | | | | | |
| Ifrd1 | | | 0.5835535 | 2.37E-12 | 0.3538238 | 3.58E-06 | 0.3390364 | 2.46E-07 | | | -0.380315 | 1.92E-08 | | | | | | | -0.219031 | 0.0020781 | | | | | | | | | | |

| | | | | | | | | | | | | | | | | | | | | | | | | | | | | | | |
|---|---|---|---|---|---|---|---|---|---|---|---|---|---|---|---|---|---|---|---|---|---|---|---|---|---|---|---|---|---|---|
| Igf1r | | | | | -0.188995 | 0.0064068 | -0.161708 | 0.0079156 | | | | | -0.421667 | 0.0010612 | | | | | | | | | | | | | | | | |
| Igfbp1 | | | 0.5178731 | 0.0000129 | 0.4216609 | 0.0000476 | | | | | | | | | | | | | | | | | | | | | | | | |
| Igfbp4 | | | | | | | | | | | -0.253677 | 0.0096731 | | | | | | | | | | | | | | | | | | |
| Igfbp5 | | | -0.359388 | 4.07E-07 | -0.502787 | 1.03E-11 | -0.44444 | 0.0000414 | | | | | | | | | | | | | | | | | | | | | | |
| Il1r1 | | | | | | | | | | | | | | | | | | | 0.3371754 | 0.004173 | | | | | | | | | | |
| Inmt | | | | | | | 0.1526664 | 5.69E-06 | | | -0.494315 | 2.66E-11 | 0.7012352 | 1.08E-26 | | | | | | | | | | | | | | | | |
| Isg15 | | | | | | | | | | | 0.5370764 | 1.25E-07 | | | | | | | | | | | | | | | | | | |
| Itgb5 | | | | | | | 0.1524219 | 0.0091988 | | | -0.18643 | 1.8E-09 | | | | | 0.4690972 | 6.57E-19 | | | | | | | | | | | | |
| Jun | | | | | -0.256691 | 8.22E-11 | | | | | -0.225498 | 2.45E-08 | -0.224287 | 0.0005116 | | | | | | | | | | | | | | | | |
| Kap | | | -0.757252 | 0.0042806 | -0.497946 | 2.88E-07 | | | | | | | | | | | | | | | | | | | | | | | | |
| Kcne1 | | | | | | | | | | | | | | | | | 0.511589 | 3.18E-06 | | | | | | | | | | | | |
| Kcnj1 | | | | | | | | | | | 0.1517246 | 4.45E-12 | | | | | 0.162849 | 0.0022544 | | | | | | | | | | | | |
| Kcnj10 | | | | | | | | | | | | | | | | | 0.151822 | 0.0039148 | 0.2601847 | 0.000057 | | | | | | | | | | |
| Kcnj16 | | | -0.118706 | 0.0056281 | -0.188085 | 6.22E-08 | | | -0.481472 | 0.0016827 | 0.49018 | 2E-87 | -0.13881 | 0.0007073 | | | 0.1809124 | 0.0014535 | 0.1634915 | 0.0047846 | | | | | | | | | | |
| Kl | | | -0.204881 | 4.79E-11 | -0.343829 | 2.96E-35 | -0.270749 | 1.33E-24 | | | 0.411103 | 1.92E-31 | | | -0.253837 | 0.0039479 | 0.5283278 | 5.78E-23 | 0.8881949 | 1.01E-10 | | | | | | | | | | |
| Klf4 | | | | | | | | | | | -0.600714 | 0.002338 | | | | | | | | | | | | | | | | | | |
| Lamc2 | | | | | | | | | 0.6368675 | 0.0000848 | | | | | | | | | -0.253497 | 0.0017157 | | | | | | | | | | |
| Lcn2 | | | | | | | | | | | 0.5109076 | 0.0008179 | | | | | | | | | | | | | | | | | 1.0648661 | 0.0010891 |
| Lgals3 | | | | | | | | | | | 0.2350548 | 0.0002637 | | | | | | | -0.198168 | 0.0006742 | | | | | | | | | 0.8231113 | 0.0036847 |
| Lpl | | | -0.350578 | 0.0014274 | -0.292863 | 0.0003237 | -0.261527 | 1E-10 | | | -0.803899 | 1.17E-160 | 0.1824964 | 0.0012504 | 0.4953214 | 1.31E-06 | 0.5036344 | 0.0021028 | | | | | | | | | | | | |
| Ly6e | | | | | | | | | | | 0.2852679 | 0.0000266 | 0.325029 | 1.23E-08 | | | 0.4140244 | 7.93E-12 | 0.3927498 | 5.01E-16 | | | 0.5685798 | 0.0083486 | 0.3430989 | 0.0027135 | 0.4557613 | 0.0001386 | | |
| Lypd2 | | | | | | | | | -1.635085 | 7.47E-07 | | | | | | | | | | | | | | | | | | | | |
| Map2k2 | | | | | | | 0.1432185 | 0.0040916 | 0.3919734 | 0.0004923 | 0.1255297 | 0.0084909 | | | | | 0.4219547 | 2.88E-07 | | | | | | | | | | | | |
| Mapk14 | | | | | -0.179778 | 3.28E-07 | -0.166726 | 6.99E-07 | | | -0.32772 | 3.15E-14 | | | | | | | | | | | | | | | | | | |
| Mapk8 | | | | | | | | | | | | | | | | | | | | | | | | | | | 1.4290422 | 0.0046959 | | |
| Mapk9 | | | | | | | | | | | -0.241235 | 0.0000176 | | | | | | | | | | | | | | | | | | |
| Mef2a | | | | | | | | | | | -0.188395 | 0.0043159 | | | | | | | | | | | | | | | | | | |
| Mir17hg | | | | | | | -0.745352 | 0.0003273 | | | | | | | | | | | | | | | | | | | | | | |
| Mt2 | | | | | | | | | | | -0.55472 | 0.0000211 | | | | | | | | | | | | | | | | | | |
| Myc | | | | | | | | | | | -0.580408 | 6.05E-06 | | | | | | | | | | | | | | | | | | |
| Napsa | | | | | -0.363842 | 0.0000011 | | | | | | | | | | | | | | | | | | | | | | | | |
| Nbl1 | | | | | | | | | -0.463613 | 0.0001649 | | | -0.756528 | 8.19E-10 | | | | | | | | | | | | | | | | |
| Nedd4l | | | | | | | | | 0.4646991 | 3.15E-06 | | | | | | | -0.306241 | 0.0000042 | -0.287283 | 2.61E-07 | | | | | | | | | | |
| Nfat5 | | | | | -0.359347 | 0.0007093 | | | | | | | -0.406045 | 0.0000112 | -0.568889 | 0.0042752 | | | | | | | | | | | | | | |
| Npnt | | | | | | | | | | | | | | | | | 0.306002 | 0.0004856 | | | | | | | | | | | | |
| Nr0b2 | | | | | | | | | | | -0.442419 | 1.38E-08 | | | | | | | | | | | | | | | | | | |
| Nr1h4 | | | 0.1793359 | 6.8E-07 | | | | | | | | | | | | | | | | | | | | | | | | | | |
| Nr3c2 | | | | | | | | | -0.876299 | 0.0069015 | | | -0.266189 | 0.0032581 | | | | | | | | | | | | | | | | |
| Nupr1 | | | | | | | | | -0.589645 | 0.007766 | -1.136781 | 8.64E-07 | | | | | | | | | | | | | | | | | | |
| Odc1 | | | | | | | -0.244662 | 3.99E-13 | -0.411056 | 6.78E-08 | -0.266776 | 9.09E-16 | | | | | | | -0.339095 | 6.22E-08 | | | | | | | | | | |
| Osgin1 | | | 0.3367424 | 4.92E-07 | 0.1661997 | 0.0017881 | | | | | | | | | | | | | 0.5925769 | 0.0001954 | | | | | | | | | | |
| Pappa2 | | | | | | | | | | | 2.054047 | 9E-09 | | | | | | | | | | | | | | | | | | |
| Pcbd1 | | | | | | | | | -0.19717 | 0.0096731 | -0.309966 | 7.71E-47 | 0.2618003 | 7.56E-15 | | | 0.4424275 | 3.01E-12 | -0.266048 | 0.0009037 | | | 0.1911168 | 0.0025587 | | | | | | |
| Pck1 | | | -0.162276 | 0.0003105 | -0.292102 | 1.22E-14 | -0.147398 | 0.0000393 | | | | | | | | | | | | | | | | | | | | | | |
| Pdgfb | | | | | | | | | | | -1.357117 | 2.13E-26 | | | | | | | | | | | | | | | | | | |
| Pdk1 | | | | | | | | | | | | | | | | | 0.439955 | 0.0002174 | 0.6737199 | 0.0000592 | | | | | | | | | | |
| Pdzk1ip1 | | | | | | | 0.1814624 | 1.22E-11 | | | | | | | | | | | | | | | 0.3539214 | 9.07E-09 | | | | | | |
| Pfkm | | | | | | | | | -0.389946 | 0.0046101 | -0.114476 | 0.0001151 | | | | | 0.2201309 | 0.0005916 | | | | | 0.2546685 | 0.0000794 | | | | | | |
| Phlda1 | | | | | | | 1.2546098 | 0.0003019 | | | | | | | | | 0.5364642 | 0.0004579 | 0.5271038 | 0.0013301 | | | | | | | | | | |
| Pik3ca | | | -0.393354 | 0.0005202 | -0.395314 | 1.73E-06 | -0.398503 | 0.0000766 | | | | | | | | | | | | | | | | | | | | | | |
| Pim3 | | | 0.2049215 | 5.56E-07 | | | 0.1599704 | 0.0001386 | | | -0.13865 | 0.0002556 | | | | | | | | | | | | | | | 0.637233 | 0.0053436 | | |
| Pkd2 | | | -0.279554 | 0.000082 | -0.225665 | 0.0000747 | -0.234374 | 0.0004923 | | | -0.204708 | 1.81E-08 | | | | | | | | | | | | | | | | | | |
| Plaur | | | | | | | | | | | | | | | | | | | -0.917094 | 1.66E-13 | | | | | | | | | | |
| Ppp1r1a | | | | | | | | | -1.546417 | 4.91E-07 | -0.057531 | 0.0039148 | | | | | | | | | | | | | | | | | | |
| Ppp3ca | | | | | | | | | -0.506721 | 0.005488 | -0.252218 | 0.0000261 | -0.310735 | 0.0017157 | | | | | | | | | | | | | | | | |
| Prkci | | | | | | | | | | | -0.190796 | 0.0005797 | | | | | | | | | | | | | | | | | | |
| Ptgds | | | | | | | | | | | -1.334323 | 1.84E-53 | | | | | | | | | | | | | | | | | | |
| Ptger3 | | | | | | | | | | | | | -0.638951 | 0.0000876 | | | | | | | | | | | | | | | | |
| Ptgs2 | | | | | | | | | | | | | | | | | | | -1.658794 | 0.0003643 | | | | | | | | | | |
| Ptpn1 | | | | | | | | | | | | | | | | | | | | | | | | | | | 0.5434945 | 0.0026872 | | |
| Rhbg | | | | | | | | | | | | | | | | | 0.165209 | 0.0002174 | | | | | 0.2732714 | 9.97E-08 | | | | | | |
| Rhcg | | | | | | | | | | | | | 0.1338522 | 0.0021028 | | | -0.165333 | 0.0002174 | | | | | | | | | | | | |
| Rheb | | | | | -0.12623 | 0.0061928 | | | | | -0.082811 | 0.0095106 | | | | | 0.2098535 | 0.0022544 | -0.184127 | 0.0081259 | | | | | | | | | | |
| Rock1 | | | | | -0.200523 | 0.0059051 | | | | | | | | | | | | | | | | | | | | | | | | |
| S100g | | | | | | | | | | | | | | | -0.304022 | 0.0073077 | 0.1675285 | 0.0022544 | | | | | | | | | | | | |
| Scnn1a | | | | | | | | | | | | | | | | | 0.2506549 | 0.0000175 | 0.3636799 | 1.4E-14 | | | | | | | | | | |
| Scnn1b | | | | | | | | | | | 1.1245044 | 0.0039705 | | | | | 0.3211335 | 9.56E-13 | | | | | | | | | | | | |
| Scnn1g | | | | | | | | | | | | | | | | | 0.1532624 | 0.0012269 | 0.2153106 | 1.15E-06 | | | | | 0.4420297 | 0.0059078 | | | | |
| Sfrp1 | | | | | | | | | | | 0.4781957 | 1.19E-10 | | | | | | | | | | | | | | | | | | |
| Sgk1 | | | | | | | | | | | -0.465061 | 0.0010668 | | | | | | | | | | | | | | | | | | |
| Sirt1 | | | | | -0.219921 | 0.0091502 | | | | | -0.250165 | 0.0003688 | -0.376289 | 0.0006429 | | | | | | | | | | | | | | | | |
| Slc12a1 | | | | | | | -0.683239 | 7.47E-06 | | | | | | | | | -0.922922 | 0.0000169 | -1.116086 | 0.0001147 | | | | | | | | | | |
| Slc13a3 | | | | | -0.230957 | 4.16E-06 | | | | | | | | | | | | | | | | | | | | | | | | |
| Slc16a10 | | | | | | | -0.250048 | 0.0062954 | | | -0.219239 | 0.0021371 | | | | | | | | | | | | | | | | | | |
| Slc22a18 | | | | | | | | | | | 0.3534093 | 0.0000416 | | | | | | | | | | | | | | | | | | |
| Slc22a6 | | | | | -0.227835 | 0.0000431 | 0.112399 | 0.0023011 | | | | | | | | | | | | | | | | | | | | | | |
| Slc22a8 | | | 0.1643134 | 6.54E-07 | | | 0.3423202 | 8.18E-28 | | | | | | | | | | | | | | | | | | | | | | |
| Slc27a2 | | | | | -0.114527 | 0.0027847 | | | | | 0.7619586 | 9.93E-29 | | | | | | | | | | | | | | | | | | |
| Slc2a1 | | | | | | | | | | | | | -0.253088 | 5.7E-08 | | | -0.225911 | 0.0002242 | -0.536908 | 1.32E-21 | | | | | | | | | | |
| Slc2a4 | | | | | | | | | | | | | 0.5882498 | 1.98E-26 | | | 0.9108631 | 1.5E-11 | | | | | 0.2283854 | 0.0007365 | 0.3924369 | 0.0020614 | | | | |
| Slc34a2 | | | | | | | | | | | 0.4607854 | 0.0000148 | | | | | | | | | | | | | | | | | | |
| Slc38a3 | | | | | | | | | | | | | | | | | 1.5380479 | 0.0091985 | | | | | | | | | | | | |
| Slc4a1 | | | | | | | | | | | | | | | | | | | | | -0.260179 | 0.0039037 | | | | | | | | |
| Slc4a3 | | | | | | | | | | | | | | | | | 0.2989264 | 0.0041555 | | | | | | | | | | | | |
| Slc4a9 | | | | | | | | | | | | | | | | | | | | | -0.295107 | 0.0049521 | | | | | | | | |
| Slc5a1 | | | | | | | 0.2474654 | 0.0001493 | | | 0.1443738 | 0.0000528 | | | | | | | | | | | | | | | | | | |
| Slc5a10 | | | 1.825098 | 2.69E-07 | | | 0.1432047 | 0.0002063 | | | | | | | | | | | | | | | | | | | | | | |
| Slc5a2 | | | | | | | -0.388639 | 1.35E-09 | | | | | | | | | | | | | | | | | | | | | | |
| Slc7a13 | | | | | | | -0.152964 | 0.0008558 | | | | | | | | | | | | | | | | | | | | | | |
| Slc8a1 | | | | | | | | | | | | | | | | | 1.4405659 | 8.67E-20 | | | | | | | | | | | | |
| Smad3 | | | | | -0.28885 | 0.0001899 | | | | | -0.229428 | 4.04E-06 | | | | | | | | | | | | | | | | | | |
| Smad7 | | | -0.2956 | 0.0004878 | -0.396005 | 5.57E-11 | -0.399342 | 7.04E-09 | | | -0.359144 | 3.45E-15 | -0.397042 | 1.95E-08 | -0.420969 | 0.0080083 | -0.377607 | 0.0000489 | -0.272422 | 0.0007126 | | | | | | | | | | |
| Socs3 | | | 0.337726 | 0.0004757 | | | 0.4636738 | 7.05E-08 | -0.412628 | 0.0000161 | -0.269512 | 2.84E-08 | | | | | | | | | | | | | | | | | | |
| Sod3 | | | | | -0.107666 | 9.78E-06 | -0.099974 | 0.0000289 | | | 1.1984175 | 2.65E-07 | | | | | | | | | | | | | | | | | | |
| Sparc | 0.3810235 | 0.0054066 | | | | | | | | | | | | | | | | | | | | | | | | | | | | |
| Sphk1 | | | | | | | | | | | -0.550721 | 2.91E-20 | | | | | | | -0.318916 | 0.0003019 | | | | | | | | | | |
| Spink8 | | | | | | | | | | | | | | | | | | | | | | | 0.2518312 | 0.000122 | | | | | | |
| Spp1 | | | | | | | | | -0.644225 | 0.0000719 | 0.4013707 | 2.64E-18 | 0.4651516 | 8.01E-10 | | | 0.3939152 | 1.12E-11 | -0.388571 | 0.0003746 | | | 0.234433 | 0.0086527 | | | | | | |
| Spp2 | | | | | 0.1645683 | 0.0002008 | 0.1944899 | 0.0009528 | | | | | | | | | | | | | | | | | | | | | | |
| Sprr1a | | | | | | | | | 1.846918 | 0.0004704 | | | | | | | | | | | | | | | | | | | | |
| Spry1 | | | | | | | | | | | | | | | | | | | 0.3709185 | 0.0086916 | | | | | | | | | | |
| St6galnac2 | | | | | -0.141186 | 0.0011924 | | | | | | | -0.208013 | 0.0060114 | | | | | | | | | | | | | | | | |
| Stat3 | | | | | | | | | 0.3037722 | 0.0056384 | | | | | | | | | -0.195054 | 0.0044191 | | | | | | | 0.354387 | 0.0039821 | | |
| Stk39 | | | | | | | | | | | | | | | | | | | | | | | 0.3378059 | 0.0016675 | | | | | | |
| Tfrc | | | 0.395401 | 3.97E-09 | 0.3226334 | 4.87E-07 | 0.3407478 | 8.23E-11 | | | -0.573322 | 8.42E-22 | 0.8045379 | 1.55E-82 | 1.189024 | 4.78E-17 | 1.4548881 | 9.88E-60 | 1.2915177 | 6.42E-55 | 0.9287456 | 0.0009954 | | | 0.8308049 | 6.11E-07 | | | | |
| Thbs1 | | | | | | | | | | | 0.3852311 | 3E-14 | -0.673747 | 7.98E-19 | -0.526957 | 0.0018071 | | | | | | | | | | | | | | |
| Timp2 | | | | | | | | | | | | | | | | | | | -0.306364 | 0.0018352 | | | | | | | | | | |

| | | | | | | | | | | | | | | | | | | | | | | | | | | | | | | |
|---|---|---|---|---|---|---|---|---|---|---|---|---|---|---|---|---|---|---|---|---|---|---|---|---|---|---|---|---|---|---|
| Timp3 | | | -0.244976 | 1.23E-07 | -0.265615 | 8.2E-13 | | | -0.688787 | 6.88E-13 | | | -0.484163 | 2.7E-10 | | | | | | | | | -1.072348 | 4.35E-08 | | | | | | |
| Tkfc | | | | | | | 0.1278449 | 0.000831 | | | 0.2840376 | 0.0085505 | | | | | | | | | | | | | | | | | | |
| Tmbim6 | | | | | -0.073535 | 0.001399 | | | | | -0.068119 | 0.0002103 | | | | | | | | | | | | | | | 0.214312 | 0.0048935 | | |
| Tmem252 | | | 0.2504919 | 7.7E-12 | | | 0.1309068 | 0.0000287 | 0.8846544 | 0.0027135 | | | | | | | | | | | | | | | | | | | | |
| Tmem33 | | | | | | | | | | | 0.1744202 | 3.54E-06 | | | | | 0.294349 | 0.0003661 | | | | | | | 0.3769584 | 0.0005064 | | | | |
| Tmigd1 | | | 0.8447434 | 0.0028013 | | | 0.2370486 | 0.0002067 | | | | | | | | | | | | | | | | | | | | | | |
| Tmsb4x | | | | | | | | | | | 0.5706908 | 2.01E-06 | | | | | | | | | | | | | | | 0.3784782 | 0.0001077 | | |
| Trf | | | 0.3864166 | 0.0005401 | 0.6039193 | 0.0000186 | | | | | | | | | | | | | | | | | | | | | | | | |
| Trp53 | | | | | | | | | 0.5062289 | 0.0003097 | 0.2246972 | 0.0004111 | | | | | | | | | | | | | 0.6083782 | 0.0002781 | | | | |
| Trpc1 | | | | | | | -0.20391 | 0.000102 | | | | | | | | | | | | | | | | | | | | | | |
| Trpm6 | | | | | | | | | | | | | | | | | 0.7091308 | 0.0004234 | | | | | | | | | | | | |
| Trpv5 | | | | | | | | | | | | | | | | | 0.2719544 | 9.67E-07 | 0.9988929 | 8.17E-06 | | | | | | | | | | |
| Ugt3a2 | | | | | -0.149517 | 0.001358 | | | | | | | | | | | | | | | | | | | | | | | | |
| Vegfa | | | | | -0.272602 | 1.1E-11 | -0.193095 | 0.0055069 | -0.800371 | 6.27E-11 | -0.66293 | 4.39E-69 | -0.236298 | 0.0000359 | | | -0.241966 | 0.0033997 | | | | | | | | | | | | |
| Wfdc2 | | | | | | | | | | | 0.5835594 | 1.84E-89 | 0.1413539 | 0.0099383 | | | 0.4043195 | 1.81E-17 | | | | | 0.3789863 | 0.0006076 | | | | | | |
| Wfs1 | | | | | | | | | | | | | | | | | 0.4236873 | 0.000831 | | | | | | | | | | | | |
| Wnk1 | | | | | | | | | | | 0.5081804 | 1.65E-26 | -0.166661 | 1.03E-07 | -0.209949 | 0.0049568 | 0.6376636 | 1.04E-09 | | | | | | | | | | | | |
| Wnk4 | | | | | | | | | | | | | | | | | 0.6762024 | 4.11E-27 | 0.5780163 | 2.01E-09 | | | | | | | | | | |
| Wnt4 | | | | | | | | | | | | | | | | | | | | | | | | | | | | | -0.656805 | 0.0020614 |
| Wnt7b | | | | | | | | | | | -0.474931 | 3.15E-11 | | | | | | | -0.532545 | 6.91E-06 | | | | | | | | | -0.807533 | 0.0005509 |
| Wnt9b | | | | | | | | | | | -1.567663 | 0.0086916 | | | | | | | | | | | | | | | | | -0.930124 | 0.0000013 |
| Xbp1 | | | -0.23567 | 9.44E-07 | -0.15884 | 0.0001386 | | | -0.247166 | 0.004127 | | | | | | | 0.4126699 | 1.1E-12 | | | | | -0.244263 | 0.0036303 | | | | | | |
| Yap1 | | | | | | | | | | | | | -0.153075 | 0.008724 | | | | | -0.307069 | 0.0000693 | | | | | | | | | | |

**Supplementary Table S1: Differentially regulated genes within cell clusters from vehicle- vs. furosemide-treated mice**

Cell clusters are labelled above. For all significantly differentially expressed genes, normalized z-score is followed by false discovery rate (FDRgray) (z-score | FDR)

**Supplementary Table S2: Absolute Cell Number by Segment / Cell Type**

| Cell Cluster | Cell Number (#) (Mean ± SEM) | | p-value | FDR |
|---|---|---|---|---|
| | Vehicle | Furosemide | | |
| Glomeruli | 83.3 ± 12.7 | 88.0 ± 15.3 | 0.828245 | 0.869943 |
| Proximal S1/S2 | 507.7 ± 53.2 | 481.0 ± 89.4 | 0.869943 | 0.869943 |
| Proximal S1/S2/S3 | 486.7 ± 13.1 | 437.7 ± 36.6 | 0.574825 | 0.722723 |
| Proximal S2/S3 | 638.3 ± 114.0 | 853.3 ± 85.2 | 0.066743 | 0.170632 |
| Thin Descending Limb | 114.7 ± 4.8 | 80.3 ± 32.9 | 0.055024 | 0.170632 |
| TAL(*) | 874.7 ± 43.4 | 499.3 ± 101.7 | 0.000587 | 0.00619 |
| DCT1 | 268.0 ± 39.8 | 365.7 ± 51.5 | 0.097369 | 0.208648 |
| DCT2(*) | 53.3 ± 10.9 | 108.3 ± 2.0 | 0.000825 | 0.00619 |
| CNT(*) | 133.3 ± 8.4 | 238.7 ± 48.9 | 0.004927 | 0.024634 |
| CCD PCs | 180.3 ± 26.1 | 228.0 ± 71.1 | 0.305724 | 0.573233 |
| Alpha ICs | 53.0 ± 9.6 | 60.3 ± 12.7 | 0.578178 | 0.722723 |
| Beta ICs | 109.3 ± 12.9 | 99.3 ± 4.6 | 0.721337 | 0.832312 |
| MCD PCs | 75.0 ± 4.0 | 49.3 ± 2.3 | 0.068253 | 0.170632 |
| Immune Cells | 99.0 ± 16.3 | 83.0 ± 17.1 | 0.372738 | 0.621229 |
| Cluster X | 15.7 ± 2.7 | 20 ± 6.5 | 0.423271 | 0.634906 |

*, indicates cell clusters with FDR < 0.1. Glom, glomerular cells; S1/S2; Proximal tubule subsegments S1/S2, S1/S2/S3, S2/S3; TAL, thick ascending limb; DCT1, distal convoluted

tubule segment 1; DCT2, distal convoluted tubule segment 2; CNT, connecting tubule; CCD PCs, principal cells of the cortical collecting duct; Alpha ICs, type A intercalated cells; Beta ICs, type B intercalated cells; MCD PCs, principal cells of the outer medullary collecting duct; Immune cells, lymphocytes and macrophages; Cluster X, *Wnt4*(+) cells. FDR, false discovery rate.

**Supplementary Table S3: Proportion of Total Cells by Segment / Cell Type**

| Cell Cluster | Proportion of Total Cells (%) (Mean ± SEM) | | p-value | FDR |
|---|---|---|---|---|
| | Vehicle | Furosemide | | |
| Glomeruli | 0.023 ± 0.003 | 0.024 ± 0.002 | 0.763894068 | 0.81845793 |
| Proximal S1/S2 | 0.139 ± 0.016 | 0.134 ± 0.029 | 0.89107905 | 0.89107905 |
| Proximal S1/S2/S3 | 0.132 ± 0.006 | 0.120 ± 0.008 | 0.222312065 | 0.416835122 |
| Proximal S2/S3(*) | 0.171 ± 0.023 | 0.232 ± 0.005 | 0.008486606 | 0.025459819 |
| Thin Descending Limb | 0.031 ± 0.001 | 0.021 ± 0.006 | 0.092634717 | 0.198502966 |
| TAL(*) | 0.237 ± 0.006 | 0.133 ± 0.012 | 1.36E-14 | 2.05E-13 |
| DCT1(*) | 0.072 ± 0.008 | 0.099 ± 0.009 | 0.020556344 | 0.051390861 |
| DCT2(*) | 0.014 ± 0.002 | 0.030 ± 0.003 | 2.13E-05 | 0.000159391 |
| CNT(*) | 0.036 ± 0.003 | 0.064 ± 0.009 | 0.002187447 | 0.010937234 |
| CCD PCs | 0.049 ± 0.007 | 0.060 ± 0.013 | 0.452156971 | 0.631270702 |
| Alpha ICs | 0.014 ± 0.002 | 0.016 ± 0.003 | 0.564832715 | 0.651730055 |
| Beta ICs | 0.029 ± 0.002 | 0.027 ± 0.002 | 0.505016562 | 0.631270702 |
| MCD PCs(*) | 0.020 ± 0.002 | 0.014 ± 0.001 | 0.002965 | 0.01112 |
| Immune Cells | 0.027 ± 0.005 | 0.023 ± 0.004 | 0.481869761 | 0.631270702 |
| Cluster X | 0.004 ± 0.001 | 0.005 ± 0.001 | 0.426804102 | 0.631270702 |

*, indicates cell clusters with FDR < 0.1. Glom, glomerular cells; S1/S2; Proximal tubule subsegments S1/S2, S1/S2/S3, S2/S3; TAL, thick ascending limb; DCT1, distal convoluted

tubule segment 1; DCT2, distal convoluted tubule segment 2; CNT, connecting tubule; CCD PCs, principal cells of the cortical collecting duct; Alpha ICs, type A intercalated cells; Beta ICs, type B intercalated cells; MCD PCs, principal cells of the outer medullary collecting duct; Immune cells, lymphocytes and macrophages; Cluster X, *Wnt4*(+) cells. FDR, false discovery rate.

**Supplementary Table S4: Primers for genotyping**

| Gene | | Sequence |
|---|---|---|
| *Igf1r*[1] | Forward | 5'-TCCCTCAGGCTTCATCCGCAA-3′ |
| | Reverse | 5'-CTTCAGCTTTGCAGGTGCACG-3' |
| Pax8-rtTA[2] | Forward | 5'-CCATGTCTAGACTGGACAAGA-3' |
| | Reverse | 5'-CTCCAGGCCACATATGATTAG'3' |
| Tet-O-Cre[2] | Forward | 5'-ATGTCCAATTTACTGACCG-3' |
| | Reverse | 5'-CGCCGCATAACCAGTGAAAC-3' |

**REFERENCES**


1. Stachelscheid, H., *et al.* Epidermal insulin/IGF-1 signalling control interfollicular morphogenesis and proliferative potential through Rac activation. *EMBO J* **27**, 2091-2101 (2008).
2. Traykova-Brauch, M., *et al.* An efficient and versatile system for acute and chronic modulation of renal tubular function in transgenic mice. *Nat Med* **14**, 979-984 (2008).
